# High-quality genome assembly of several *Noccaea caerulescens* accessions reveals gene expansion events associated with metal adaptation and hyperaccumulation

**DOI:** 10.64898/2026.10.20.760424

**Authors:** Maria Kuzmik, Ophélie Courtin, Christophe Klopp, Marie-Christine Carpentier, Nathalie Rodde, Caroline Callot, Anne Ausset, Céline Roques, Céline Vandecasteele, Véronique Gautier, Elodie Belmonte, Elodie Flaven, Céline Devaux, Agnès Mignot, Marie Mirouze, Sylvain Merlot

## Abstract

Since the beginning of the 20th century, the growth of the world’s population and industrial development have led to increased soil pollution with metals, which poses a threat to the environment and human health. In this context, plants that are adapted to metalliferous soils and hyperaccumulate metals offer an opportunity to develop phytotechnologies for remediating metal-polluted soils. However, the molecular mechanisms by which plants tolerate and hyperaccumulate metals are not yet fully understood, which limits the improvement of these technologies. The metal hyperaccumulator species *Noccaea caerulescens*, belonging to the Brassicaceae family, is being developed as a genetic model to decipher these mechanisms. In this study, we have significantly improved the genome assembly of the *N. caerulescens* Cira accession and have produced high-quality assemblies for five additional accessions originating from different soil types and genetic groups. We used these high-quality assemblies to identify gene copy number variation associated with the metal hyperaccumulation trait in the Noccaea genus. Our analysis revealed that gene copy number variation affects a wide variety of biological functions in *Noccaea*, including genes that encode metal transporters belonging to the HMA, NRAMP and ZIP families. Notably, duplications affecting the magnesium transporter *MGR2* suggest that the adaptation to ultramafic soils may be an ancestral trait in *Noccaea*. Finally, the association of transposable elements with gene duplications suggests that transposon activity may have played an important role in the evolution of metal hyperaccumulation in this genus.

## 1. Introduction

Since the beginning of the 20th century, the growth of world’s population and industrial development have led to profound changes that are proposed to mark a new geological epoch called the Anthropocene. These changes include increased soil pollution with metals resulting from the burning of fossil fuels, mining activities, and the use of pesticide and fertilizer in agriculture (Hou et al., 2025, 2020; Wagreich and Draganits, 2018; Waters et al., 2016). The current development of low-carbon energy, which relies on the extensive use of metals such as copper, nickel, and cobalt, could further increase the risk of environment metal pollution (Lèbre et al., 2020; Sonter et al., 2020; Vidal et al., 2022). In this context, plant species that are adapted to metalliferous soils and hyperaccumulate metals provide an opportunity to develop phytotechnologies for extracting metals from contaminated soils and recycle them from the biomass (Huang et al., 2026; Rylott and van der Ent, 2025; van der Ent et al., 2026). As with traditional crop species, developing these phytotechnologies would greatly benefit from a better understanding of the molecular mechanisms involved in metal tolerance and hyperaccumulation.

Over the past 20 years, several genetic resources and transformation protocols have been developed to establish *Noccaea caerulescens* as a genetic model to decipher the molecular mechanisms of metal tolerance and hyperaccumulation in plants (Assunção et al., 2003b; Blande et al., 2017; Milner and Kochian, 2008; Wang et al., 2022; Yamjabok et al., 2026). *N. caerulescens* is a nickel, zinc and cadmium hyperaccumulator of the Brassicaceae family. It belongs to the Coluteocarpea tribe, which includes other metal hyperaccumulators of the *Noccaea* genus, such as *N. praecox* (Jakovljević et al., 2024; Vogel-Mikuš et al., 2005), and with non-accumulator species of the *Microthlaspi* genus, including *M. erraticum* and *M. perfoliatum* (Ali et al., 2016; Hendriks et al., 2023; Mishra et al., 2020). However, some *M. perfoliatum* populations from Turkey could have acquired the capacity to accumulate nickel (Çelik et al., 2018; Reeves et al., 2008). *N. caerulescens* grows naturally on both non-metalliferous and metalliferous soils including ultramafic soils - also known as serpentine - which are rich in nickel, as well as calamine soils, which are rich in zinc and cadmium. More than 60 accessions, which belong to at least three genetic units originating from historical events, and hyperaccumulating different levels of nickel, zinc and cadmium, depending on the type of soil they are growing on, have been identified in Western Europe (Assunção et al., 2003; Escarré et al., 2000; Gonneau et al., 2017, 2014; Sterckeman et al., 2017). Ex-situ experiments further indicated that zinc hyperaccumulation capacity is constitutive in the species, while the capacity to hyperaccumulate nickel and cadmium is more variable (Kozhevnikova et al., 2020; Sterckeman et al., 2017). In particular, accessions originating from the southern part of the Massif Central region in France, including Ganges, display the highest capacity to hyperaccumulate cadmium (Sterckeman et al., 2017).

From a genetic point of view, *N. caerulescens* is a diploid species and has a mixed mating system, characterized by variable amounts of self-fertilization. Its nuclear genome consists of seven chromosomes originating from the Proto-Calepinaea karyotype, totaling approximately 270 Mb (Mandáková et al., 2015; Mandáková and Lysak, 2008). Sequencing and assembling the *N. caerulescens* genome has been challenging because short-read sequencing technologies could not resolve long stretches of repetitive sequences, which led to highly fragmented assemblies (Kiefer et al., 2019). Recently, however, a high-quality genome assembly of the Cira accession totaling 254 Mb was obtained using PacBio HiFi long-read sequencing technology and Hi-C scaffolding (Belloeil et al., 2026). While this assembly represents an important improvement of the *N. caerulescens* genome assembly, it also reveals discrepancies with the structure of the genome inferred from cytogenetic analysis (Mandáková et al., 2015). Intra- and interspecific comparative transcriptomic analyses have identified genes whose expression is associated with metal hyperaccumulation in *N. caerulescens* (Belloeil et al., 2026; García de la Torre et al., 2021; Halimaa et al., 2014; Hammond et al., 2006; van de Mortel et al., 2006). These analyses revealed that metal hyperaccumulation is associated with high expression of genes involved in the metal homeostasis regulation. This includes several metal transporters from the heavy metal ATPase (HMA), ZRT and IRT-like proteins (ZIP), natural resistance-associated macrophage proteins (NRAMP), and iron regulated/Ferroportin (IREG/FPN) families. The NcHMA4 transporter can export zinc and cadmium out of plant cells (Bernard et al., 2004; Craciun et al., 2012; Leitenmaier et al., 2011; Papoyan and Kochian, 2004). Similar to its orthologs in *Arabidopsis*, it has been proposed that the physiological function of NcHMA4 is to load zinc and cadmium into the xylem for translocation to the shoot (Hanikenne and Nouet, 2011). Conversely, NcHMA3 is localized in vacuolar membrane, and is likely to play a role in cadmium storage in mesophyll and epidermal cells of leaves (Ueno et al., 2011). The transporter NcIREG2 was shown to play a similar role for the sequestration of nickel in root and leaf cell vacuoles. More generally, the high activity of vacuolar IREG transporters in leaves was proposed to correspond to a highly convergent mechanisms involved in the evolution of nickel hyperaccumulation in plants (García de la Torre et al., 2021; González et al., 2024; Meier et al., 2018; Merlot et al., 2014). The orthologue of the iron-uptake transporter IRT1 is constitutively expressed in roots of *N. caerulescens*, and mutations of NcIRT1 in several accessions revealed its role in the uptake and accumulation of both cadmium and nickel in this species (Belloeil et al., 2026; Halimaa et al., 2019). In addition to NcIRT1, the plasma membrane transporter NcNRAMP1 was proposed to mediate cadmium uptake and radial transport to the stele (Milner et al., 2014).

Therefore, identifying the mechanisms responsible for the high expression of metal homeostasis genes is critical to understand the evolution of metal hyperaccumulation in *N. caerulescens*. In another metal hyperaccumulator of the Brassicaceae family, *Arabidopsis halleri*, the high expression of *AhMHA4* has been shown to result from tandem triplication and modifications of cis-regulatory elements (Castanedo et al., 2025; Hanikenne et al., 2013, 2008). Duplication of genes encoding the HMA4, HMA3, and NRAMP1 transporters has also been observed in several *N. caerulescens* accessions (Belloeil et al., 2026; Craciun et al., 2012; Milner et al., 2014; Ó Lochlainn et al., 2011; Ueno et al., 2011). These findings suggest that variations in gene copy number (CNV) play a role in the high expression of these metal transporters, and the evolution of metal hyperaccumulation in *N. caerulescens*. However, it is still unclear if the high expression of these genes also results from the modification of cis-regulatory elements and if other loci involved in metal hyperaccumulation are affected by CNV in *N. caerulescens*.

In this study, we improved the assembly of the *N. caerulescens* Cira genome by using ultra-long DNA reads. We also generated new, high-quality assemblies from additional *N. caerulescens* accessions originating from different edaphic and genetic groups. These high-quality assemblies facilitated the structural annotation of genes and transposable elements (TE) in this species. Using these data, we performed comparative genomic analysis with non-accumulator species of the Brassicaceae family to identify gene expansion associated with metal hyperaccumulation in *Noccaea* species. Our findings suggest that TE activity has played an important role in the duplication of genes encoding metal transporters, and therefore in the evolution of metal hyperaccumulation in *N. caerulescens*.

## 2. Materials and methods

### 2.1. Plant material

The collection of *Noccaea caerulescens* accessions Cira (*Ncci*), Firmiensis (*aka* Puy de Wolf, *Ncfi*) and Bergenbach (*Ncbe*) seeds was previously described (Belloeil et al., 2026). Seeds of the accessions Monte Prinzera (*Ncmp*) and Col de Marchairuz (*Ncmar*) were obtained from the laboratory of Mark Aarts (Wageningen University and Research, The Netherland). Plants were grown from seeds in hydroponics for eight weeks, as previously described (Belloeil et al., 2026). For each accession, young leaves were collected from a single plant and frozen in liquid nitrogen before DNA extraction. Leaves of the Saint-Laurent-Le-Minier accession (*Ncslm*) were collected directly from a single plant at Les Avinières tailing pond (43°55’59.3"N 3°40’00.9"E).

### 2.2. Genomic data

For comparative genomic analyses, we used reference genome assemblies from *Noccaea praecox* (GCA_051546285.1), *Microthlaspi erraticum* (GCA_902728155.2), *Thlaspi arvense* (GCA_911865555.2), *Arabidopsis thaliana* (GCA_000001735.2), *Eutrema salsugineum* (GCA_000478725.1), and *Carica papaya* (GCA_000150535.1).

For EGAPx annotation, we used the RNA-seq data from *Noccaea caerulescens* (SRX24619841, SRX24619844) *Microthlaspi perfoliatum* (SRX24730396, SRX24730395), *Thlaspi arvense* (SRR29820641, SRR29820626), *Eutrema salsugineum* (SRR30508955, SRR30508947), and *Carica papaya* (SRR7145705, SRR7145703).

### 2.3. DNA extraction and sequencing

For *Ncci*, high molecular weight genomic DNA was extracted from 6g of young, frozen leaves using Qiagen Genomic-tip G500 columns, essentially according to the manufacturer’s instructions (Supplemental methods). The DNA (20 µg) was then sized from 5 to 100 kb using a SageELF instrument (Sage Science) according to manufacturer’s instruction and purified using AMPure beads (Beckman Coulter). The *Ncci* DNA library was prepared using the Oxford Nanopore Technology (ONT) Ligation Sequencing Kit v14, following the manufacturer protocol with minor changes (Supplemental methods). The library was sequenced on an ONT MinIon flow-cell (FLO-MIN114) following the manufacturer instructions. The data were basecalled from pod5 files with the super-accurate model using the Dorado tool (ONT, v0.7.3). In total, 5.3 GB of raw data were generated with a N50 of 38 kb (SRR32973374).

For *Ncslm,* high molecular weight DNA was extracted from 500 mg of leaves using an SDS/CTAB based protocol and purified with magnetic beads on a KingFisher Flex system. (Supplemental methods). A SMRTBell library was prepared from 10 µg DNA sheared at 25 kb (Supplemental methods) and sequenced on a PacBio Sequel II instrument using the CCS mode. A total of 26.2 Gb of sequence (N50 24 kb) was generated (SRR33000239)

For the other *N. caerulescens* accessions, *Ncfi*, *Ncbe*, *Ncmp* and *Ncmar*, frozen leaves (1.5 g) were grinded with pestle and mortar and genomic DNA extracted with a NucleoBond HMW DNA extraction kit (Macherey-Nagel) as previously described (Belloeil et al., 2026). The genomic DNA of the accessions were multiplexed with Native Barcoding kit SQK-NBD114-24 and sequenced on a ONT PromethION A100-series machine using a flowcell type FLO-PRO114M (Supplemental methods). Sequences were basecalled with Dorado-basecall-server v7.3.11 (Super-accurate basecalling, 400 bps) from the raw pod5 file. We obtained a mean volume of 25 Gb (N50 22 kb) of data from basecalled reads per DNA sample (SRR33000240, SRR33000241, SRR33000242, SRR33000243).

### 2.4. Genome assembly, scaffolding and quality analyses

For the genome assembly of *Ncci*, the ONT long reads generated in this study (17X coverage) were combined with the previously generated *Ncci* PacBio HiFi reads (SRX24805471), reduced to a 30X coverage (Belloeil et al., 2026). These combined reads were assembled using Hifiasm v0.19.9 (Cheng et al., 2021). The contigs were scaffolded with RagTag v2.1.0 using the *Ncci* v2.3 assembly (GCA_046127275.1) as a template (Alonge et al., 2022). We verified the structure of the newly proposed scaffolds using a Hi-C contact map generated with *Ncci* OMNI-C data (SRX24805472) with Juicebox v2.15 (Durand et al., 2016). We oriented the 7 largest contigs, which correspond to the *N. caerulescens* chromosomes, according to the previously proposed *N. caerulescens* chromosome orientation (Mandáková et al., 2015). This novel assembly is called *N. caerulescens* Cira v3.1 (*Ncci* v3.1).

For the genome assembly of *Ncslm*, the complexity of the *Ncslm* PacBio HiFi reads set was reduced to a genome coverage of 35X (10 Gb) and reads assembled with Hifiasm v0.19.9. The contigs were then scaffolded with RagTag v2.1.0 using the *Ncci* v3.1 assembly as a template to generate the *Ncslm* v2.1 assembly.

For the genome assembly of *Ncfi*, *Ncbe, Ncmp* and *Ncmar*, the ONT basecalled reads were first corrected using Pecat v0.0.3 (Nie et al., 2024)and then assembled (75X coverage) with Hifiasm v0.19.9. For each accession, the contigs were scaffolded with RagTag v2.1.0 using the *Ncci* v3.1 assembly as template to generate the *Ncfi* v1.1, *Ncbe* v1.1, *Ncmp* v1.1 and *Ncmar* v1.1 assemblies.

For all accessions and before assembly, a k-mer analysis of ONT and PacBio Hifi raw reads was performed with Jellyfish v2.3 and visualized with GenomeScope v2.0 to estimate genome size, heterozygosity and genome coverage (Marçais and Kingsford, 2011; Ranallo-Benavidez et al., 2020). The consensus quality values (QV) and the k-mer completeness of the assemblies were estimated with Merqury v1.3 (Rhie et al., 2020), and D-GENIES v1.5.0 was used to evaluate large-scale structural integrity, potential misassemblies and assembly duplications (Cabanettes and Klopp, 2018). Assembly duplications were purged manually using D-GENIES and Mercury output (Supplemental Fig. 1A). Taxonomic assignment of contigs was performed using BlobTools v4.4.2 to detect potential genome contamination (Challis et al., 2020). Assembly contiguity and accuracy metrics including N50, L50, total assembly size, and the number of contigs, were computed with the assemblathon_stats.pl script found at (https://github.com/KorfLab/Assemblathon) (Bradnam et al., 2013). Genome completeness was further evaluated using BUSCO v5.7.1 with the Embryophyta lineage dataset odb10 to assess the presence of conserved single-copy orthologs (Manni et al., 2021). OMArK v0.3 provided additional gene-space and structural validation (Nevers et al., 2025).

Tandem repeat regions, characteristic of centromere regions, were identified on the seven largest contigs of the assemblies using RepeatOBserver v1 (Elphinstone et al., 2025). A Python script was used to identify the telomere regions by calculating the number of the canonical plant telomeric repeat, TTTAGGG, on both strands of DNA within a 10-kb window. The 24 genomic blocks of the ancestral Brassicaceae karyotype (ACK) were annotated on the seven largest contigs of each assembly by associating *N. caerulescens* genes with ACK block specific gene sets from *Schrenkiella parvula* using NCBI BLAST+ v2.2.28 best reciprocal hits (Murat et al., 2015). The identification of contigs corresponding to *N. caerulescens* chloroplast and mitochondrial genomes is described in Supplemental methods.

### 2.5. Structural and functional annotation of genes

To avoid biases associated with the method used for the annotation of genes in different species, all genome assemblies used in this study (except *A. thaliana*) were annotated using NCBI EGAPx v0.3.2-alpha (https://github.com/ncbi/egapx). EGAPx was run using the corresponding raw genomic sequence data, the appropriate NCBI taxonomy identifier (taxid), and when available paired-end RNA-Seq data generated from leaf and root tissues as evidence to support gene model prediction. The gene model of TAIR10 annotation was used for *A. thaliana*. The *N. caerulescens* genome assemblies generated in this study were also annotated by an alternative method. The structural annotation of the genes from the *Ncci* v2.3 assembly (Belloeil et al. 2026), was transferred to the new genome assemblies using Liftoff v1.6.3 (Shumate and Salzberg, 2024). In addition, *de novo* annotations were obtained with Helixer v0.3.1 (Stiehler et al., 2021). Both annotations were then merged using AGAT v1.2.0 (Dainat et al., 2023). The correspondence between annotated gene in *Ncci* v2.3 and *Ncci* v3.1 assemblies is given in Supplemental Table 1.

The function of genes from *N. caerulescens* accessions and the other species used in this study was annotated using the MapMan hierarchical bin system with Mercator4 v7 (Bolger et al., 2021). For *N. caerulescens* accessions, the functional description of each gene was transferred to the GFF file of the gene model annotation using a custom Python script.

### 2.6. Annotation of transposable elements

We annotated transposable elements (TEs) on the *Ncci* v3.1 genome assembly using a previously described method (Kim et al., 2024). Additionally, miniature inverted-repeat transposable elements (MITEs) were identified using MITE-Hunter (https://github.com/jburnette/MITE-Hunter), with the default parameters (Han and Wessler, 2010). These analyses resulted in the characterization of 395 non-redundant TE families, including 91 Gypsy LTRs, 38 Copia LTRs, 37 PaRa and P elements, 20 LINEs, 10 MuDRs, nine Harbingers, five hATs and helitrons, one Mariner and 142 MITEs. The annotations of TEs in the genome assemblies of other *N. caerulescens* accessions as well as in *N. praecox* and *M. erraticum* were performed with the *Ncci* v3.1 TE database using RepeatMasker v4.2.3 with the default parameters and the WUBLAST software (Smit et al., 2015).

### 2.7. Visualization of genome assemblies and synteny

We used the Circos v2.0 software for the graphical representation of the structure and annotations of the *N. caerulescens* genome assemblies (Cui et al., 2020).

Genome-wide synteny and structural genome rearrangements analysis were performed using SyRI v1.6.3 (Goel et al., 2019). Whole-genome pairwise alignments were generated using minimap2 v2.28 with the *asm5* preset, optimized for alignment of closely related genome assemblies (Li, 2018). The resulting alignments were converted to BAM format, sorted, and indexed using SAMtools v1.21 (Li et al., 2009). Synteny plots were generated with the SyRI output files using plotsr.

### 2.8. Analysis of gene copy number variations

As previously described, we used the gene models provided by EGAPx and annotated with Mercator4 v7.0 for all species but *Arabidopsis thaliana*. For *A. thaliana*, we used the Araport11 annotation directly from the Mercator4 v7.0 database. The distribution of genes from the different genomes into MapMan’s hierarchical functional bins is provided in Supplemental Data Set 1. For the comparative genomic analysis, the species were then classified as either hyperaccumulators (*i.e. N. caerulescens* accessions and *N. praecox*) or non-accumulators (all other species). We assessed differences in gene copy number, contained in lower-level bins, between the two groups using a Kruskal-Wallis test with a significance threshold of p < 0.01. Results are displayed in a heatmap generated the R package ggplot2. Barplots representing the variation in gene copy number variation (CNV) were generated using the R packages ggplot2, tidyverse and stringr (Wickham, 2016). Gene orthology among Brassicaceae species was defined by reciprocal best blast hits and gene synteny using the Plaza Dicots 5.0 framework (van Bel et al., 2022).

Pairwise sequence alignments were generated using BLASTn in CLC Genomics Workbench v25 (Qiagen) and saved as a CSV file. Coordinates of gene models and TE were extracted from the GFF files of annotated genome assemblies and exported as a Genbank file. The graphs were generated using the R packages ggplot2, tidyverse, gggenes and viridis.

### 2.7 Access to scripts and genomic data

The Python scripts used in this study can be found at https://github.com/courtin-ophelie The raw sequencing reads and assembled genomes generated in this work are available in NCBI’s BioProject PRJNA1096268. New *N. caerulescens* genome assemblies and their annotations were deposited in the recherche.data.gouv repository (doi:10.57745/OIP19T)

## 3. Results

### 3.1 An improved assembly of the Noccaea caerulescens Cira genome

Comparative genomic analyses rely on the high-quality genome assemblies of various plant species or accessions. We recently published a chromosome-level assembly of the *Noccaea caerulescens* Cira genome (*Ncci* v2.3) built with PacBio HiFi and Hi-C reads (Belloeil et al., 2026). However, the presence of long stretches of repeated sequences (>10 kb) in the *N. caerulescens* genome did not enable to resolve certain regions. To improve the assembly of the *N. caerulescens* Cira genome, we generated longer reads from the same accession using Oxford Nanopore Technologies (ONT). Hybrid assembly of the PacBio HiFi and ONT data produced 296 contigs, eight of which exceeded 10 Mb. The total assembly size was 270 Mb, with an N50 of 31.1 Mb (Table 1). A comparison with the *Ncci* v2.3 assembly revealed that single contigs fully covered chromosomes NC1, NC3, and NC7 (Fig. 1A). We reconstructed the sequences of the other chromosomes by scaffolding the remaining contigs using the *Ncci* v2.3 assembly as a reference. We then oriented the seven chromosomes according to the previously proposed structure of the *N. caerulescens* genome (Mandáková et al., 2015; Mandáková and Lysak, 2008). In addition to these chromosomes totaling 256 Mb, we identified contigs corresponding to the chloroplast and mitochondrial genomes through homology with the *Thlaspi arvense* chloroplast genome and the *Microthlaspi perfoliatum* mitochondrial genome, respectively (Supplemental Fig. 2). This novel assembly is called *Ncci* v3.1.

**Fig. 1:**
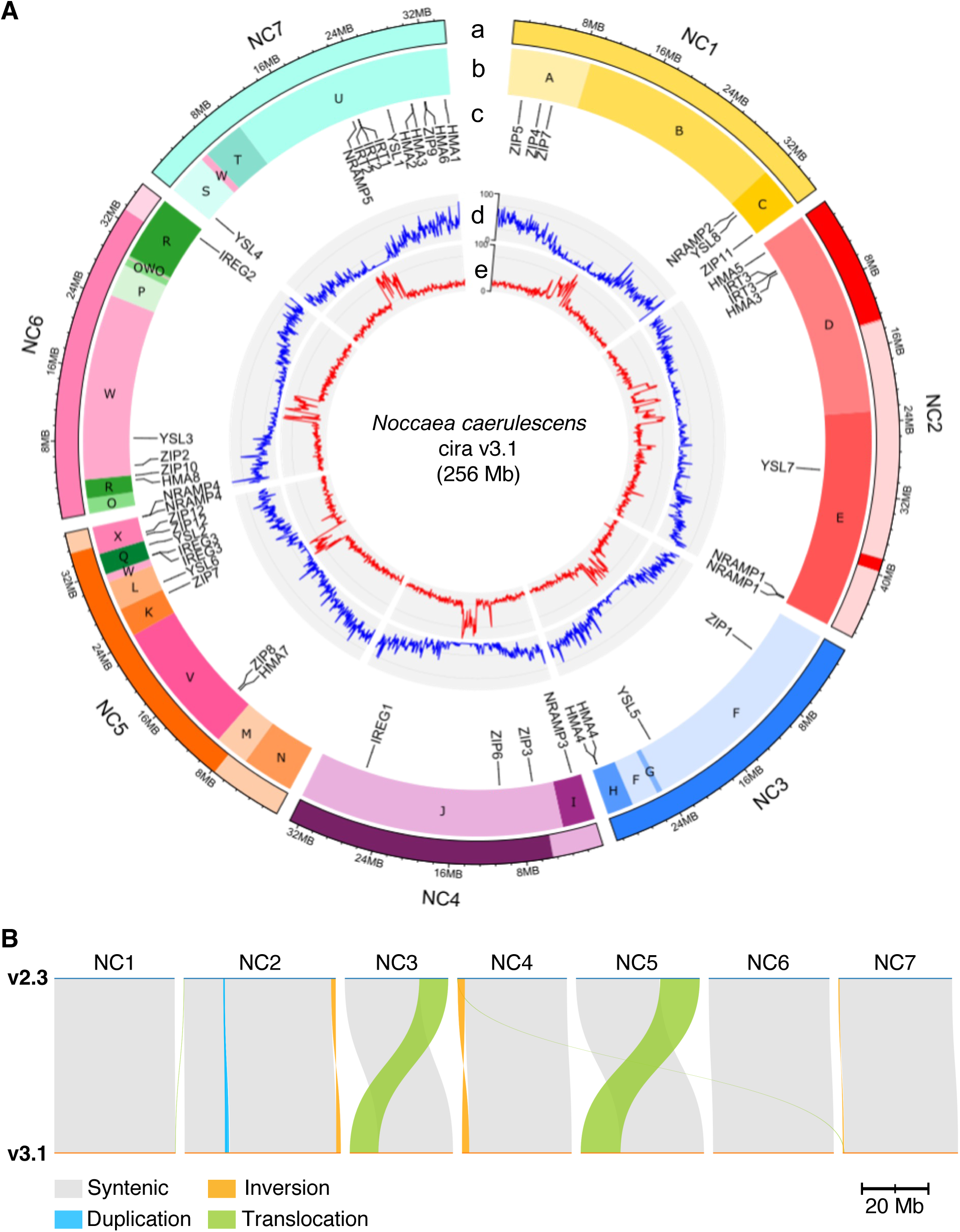
Representation of the *Noccaea caerulescens* genome assembly v3.1. (A) Representation of the seven chromosomes (NC1 to NC7) of the *N. caerulescens* Cira genome assembly v3.1 (*Ncci* v3.1). From the outermost to the innermost rings, the tracks represent the limits of the contigs forming each chromosome (a), the annotation of the ancestral Brassicaceae genomic blocks (b), the positions of marker genes coding for metal transporters annotated by EGAPx (c), the density of coding sequences (d), and tandem repeat regions calculated in 10kb windows (e). (B) Comparison of the genome assembly *Ncci* v3.1 with the previously published *Ncci* v2.3 assembly (Belloeil et al., 2026). Syntenic, duplicated, inverted and translocated regions are represented. The scale bar corresponds to 20 Mb.

**Table 1:**
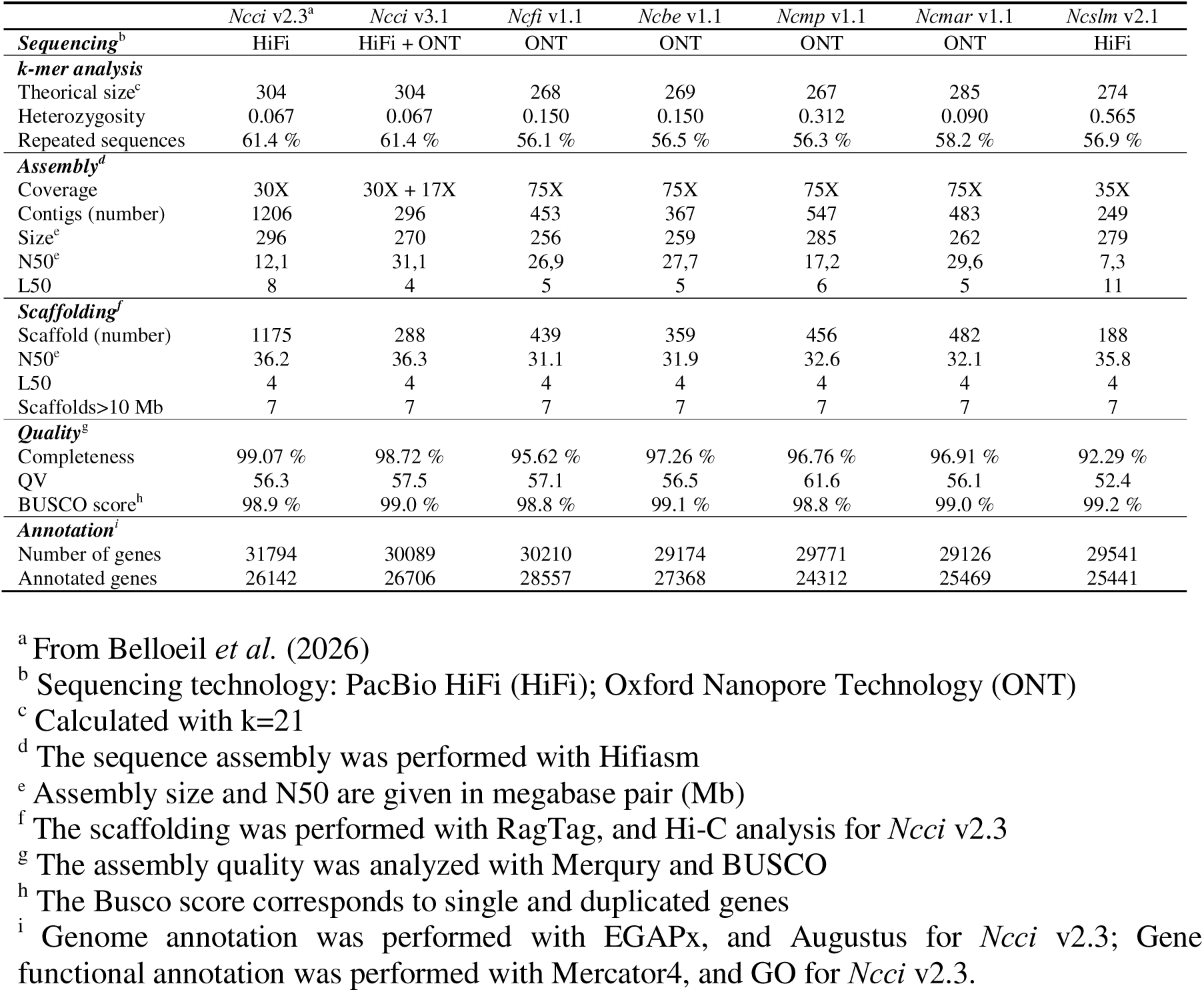
Analysis of the genome assemblies of six *Noccaea caerulescens* accessions.

Synteny analysis between the *Ncci* v2.3 and v3.1 assemblies revealed two large intrachromosomal translocations in chromosomes NC3 and NC5, as well as a few intra- and inter-chromosomal translocations and inversions of short regions (Fig. 1B; Supplemental Fig. 1C). This novel structure is supported by a Hi-C map analysis (Supplemental Fig. 1B) and by the distribution of the 24 genomic blocks of the ancestral Brassicaceae karyotype (ACK) across the seven *N. caerulescens* chromosomes (Fig. 1A). This distribution aligns with the one previously proposed for this species (Mandáková et al., 2015). We detected well defined tandem repeated regions, indicative of centromere regions, in the central region of each proposed chromosome, except for NC2 in which the tandem repeated region is more extended (Fig. 1A). These observations indicate that the *N. caerulescens* chromosomes are essentially metacentric. Finally, we detected the presence of 100 to 200 copies of the canonical plant telomeric sequence, TTTAGGG, at the extremities of all chromosomes except at the farthest end of chromosome NC5, indicating a very good completeness of the assembly in the telomeric regions (data not shown). The analysis of the contigs and scaffolds of the *Ncci* v3.1 assembly using Merqury revealed a high k-mer completeness (98.72%) and a consensus quality score (QV) of 57.5 (Table 1; Supplemental Fig. 1D). The BUSCO analysis showed that 99.0% of the 1614 core conserved genes are present as single or duplicate genes similar to the previous *Ncci* v2.3 version of the assembly (Table 1). We annotated the genomic features on the *Ncci* v3.1 assembly using EGAPx. This led to the identification of 30 089 genes, 26 706 of which (89%) received a functional annotation with Mercator 4 (Table 1). The annotation of transposable elements (TE) revealed that the *Ncci* v3.1 assembly contains 91.3 Mb of TE DNA representing 34% of the genome assembly (Supplemental Fig.3). Together, these data indicate that this novel assembly (*Ncci* v3.1) represent an improvement of the *N. caerulescens* genome assembly compared to the previously published *Ncci* v2.3 assembly (Belloeil et al., 2026).

### 3.2 Comparative analysis of N. caerulescens genome assemblies

To better understand the genomic diversity within the *N. caerulescens* species, we generated five additional genome assemblies from accessions originating from the genetic units, SU1 (Saint Laurent le Minier *Ncslm*), SU2 (Firmiensis *Ncfi*), SU3 (Bergenbach *Ncbe*, Col de Marchairuz *Ncmar*), and adapted to ultramafic (Monte Prinzera *Ncmp*, *Ncfi*, *Ncbe*), calamine (*Ncslm*) and non-metalliferous soils (*Ncmar*). The k-mer analysis of the PacBio HiFi (*Ncslm*) and the corrected ONT reads (*Ncfi*, *Ncbe*, *Ncmp*, *Ncmar*) predicted genome sizes ranging from 267 Mb to 285 Mb. We obtained assemblies composed of 249 to 547 contigs, with total assembly length ranging from 256 Mb to 278 Mb (Table1). The assemblies obtained with ONT reads contain contigs with a N50 ranging from 17.2 to 29.6 Mb, while the *Ncslm* assembly generated with PacBio HiFi reads contains contigs with a N50 of 7.3 Mb. For each accession, the contigs were scaffolded into chromosomes using the *Ncci* v3.1 assembly as reference (Supplemental Fig. 4). The analysis of these assemblies using Merqury revealed a high k-mer completeness (from 92.29% to 97.26%) and a consensus quality score (QV) ranging from 52.4 to 61.6. BUSCO analysis indicated that 98.8% to 99,2% of the 1 614 core conserved genes are present as single or duplicate genes in these assemblies. The annotation of TEs revealed that they represent from 72.2 to 92.6 Mb in these genomes, with a similar distribution of families as observed in *Ncci* (Supplemental Fig. 3). These assemblies contain from 29 126 to 30 210 protein-coding genes, 82 to 95% of which were functionally annotated (Table 1). The comparative analysis of the chromosome assemblies indicates that *Ncci* and *Ncslm* chromosomes have an excess of 20 to 30 Mb compared to the other accessions, primarily due to the increased size of chromosomes NC1, NC2 and NC7 (Fig. 2). The synteny analysis revealed a relatively conserved overall genome structure, with most structural variations located in the centromeric regions (Fig. 2; Supplemental Fig. 5). However, this analysis also revealed that the *Ncci* genome contains a large inversion in the lower arm of chromosome NC2 compared to other accessions, and that the *Ncmar* genome contains an inversion in the upper arm of chromosome NC5 (Fig. 2).

**Fig. 2:**
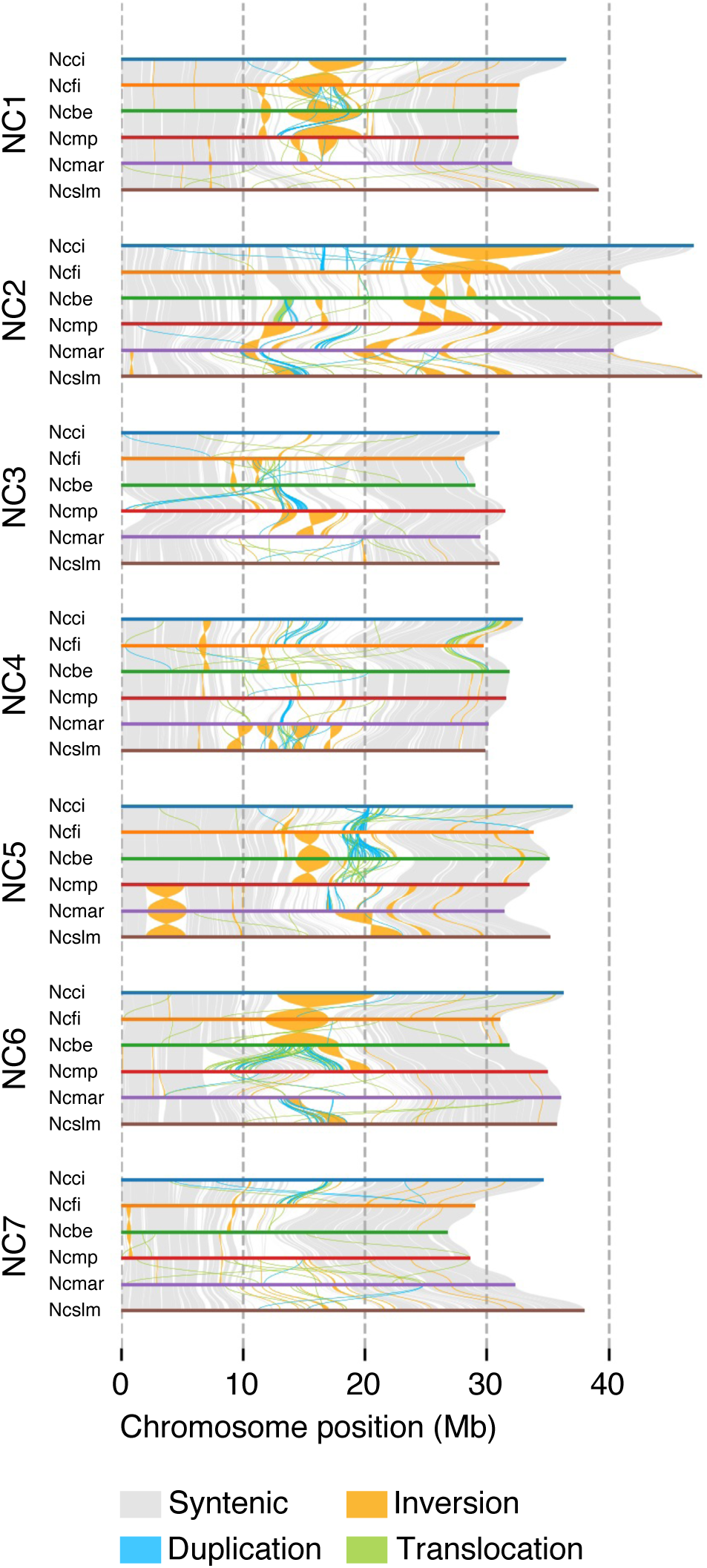
Comparison of *Noccaea caerulescens* accessions genome assemblies. The genome assemblies of the six *N. caerulescens* accessions described in this study *Ncci* v3.1, *Ncfi* v1.1, *Ncbe* v1.1, *Ncmp* v1.1, *Ncmar* v1.1, *Ncslm* v2.1 were compared according to chromosome numbers. Syntenic regions and structural rearrangements, including duplications, inversions and translocations are indicated.

These data indicate that the quality of these genome assemblies is similar to that of *Ncci* v3.1 and therefore represent a unique set of data for comparative genomics studies in *N. caerulescens* and in the Brassicaceae family.

### 3.3. Biological functions affected by gene copy number variation in the Noccaea genus

Gene copy number variation (CNV) has been associated with several adaptive traits in plants, including metal hyperaccumulation (Hanikenne et al., 2008; Ó Lochlainn et al., 2011). To identify biological functions showing gene CNV associated with metalliferous soil adaptation and metal hyperaccumulation in *the Noccaea* genus, we examined the metal hyperaccumulator group, which includes the *N. caerulescens* accessions and *Noccaea praecox*, as well as the non-accumulator group, which includes the Brassicaceae species *Eutrema salsugineum*, *Thlaspi arvense*, *Arabidopsis thaliana*, as well as *Microthlaspi erraticum* of the Coluteocarpeae tribe, to which the *Noccaea* species belong. *Carica papaya* of the order Brassicales was used as an outgroup. We classified the annotated genes from the genome of these species using the bin system of MapMan, which contains 31 top-level and 6 622 sub-level functional bins (Schwacke et al., 2019). Then, we compared the number of genes in different sub-bins between hyperaccumulator and non-accumulator species (Fig. 3). We identified 146 functional sub-bins out of 6 622 (2.2%), distributed among 27 top-level bins (out of 31) that show significant variation in gene number (Fig. 3). This result suggests that, in the *Noccaea* genus, gene CNV affects a relatively small number of genes involved in a large diversity of biological functions. 115 sub-bins (27 bins) contain more genes in the hyperaccumulator groups, while only 32 sub-bins (18 bins) contain more genes in the non-accumulator groups. We examined more specifically bins 24 and 25, containing genes involved in solute transport and nutrient uptake, because these functions are likely important for metal hyperaccumulation (Fig. 4). Six sub-bins, corresponding to genes encoding V-type proton ATPase catalytic subunits A (VHA-A, sub-bin 24.1.1.2.1), P1B-type heavy metal cation-transporting ATPases (HMA, sub-bin 24.1.2.1.1), magnesium cation transporters (MGR, sub-bin 24.2.25), NRAMP transporters (sub-bins 24.2.3.10 and 25.4.4.13.2), and type-3 metallothioneins (MT3, sub-bin 25.5.6.2), contain more genes in the metal hyperaccumulator group than in the non-accumulator group. Conversely, three sub-bins containing genes that encode the calcium/manganese cation transporters PML4/5 (sub-bin 24.2.14.4), the ATP-ADP antiporters NTT1/2 (sub-bin 24.2.2.12) and the outer membrane porins OEP24 (sub-bin 24.4.4), contain fewer genes in the hyperaccumulator group. Among *N. caerulescens* accessions, *Ncslm* display a strong increase of genes encoding MT3, HMA, and NRAMP transporters. The *Ncfi* accession displays a strong increase of genes encoding MGR transporters.

**Fig. 3:**
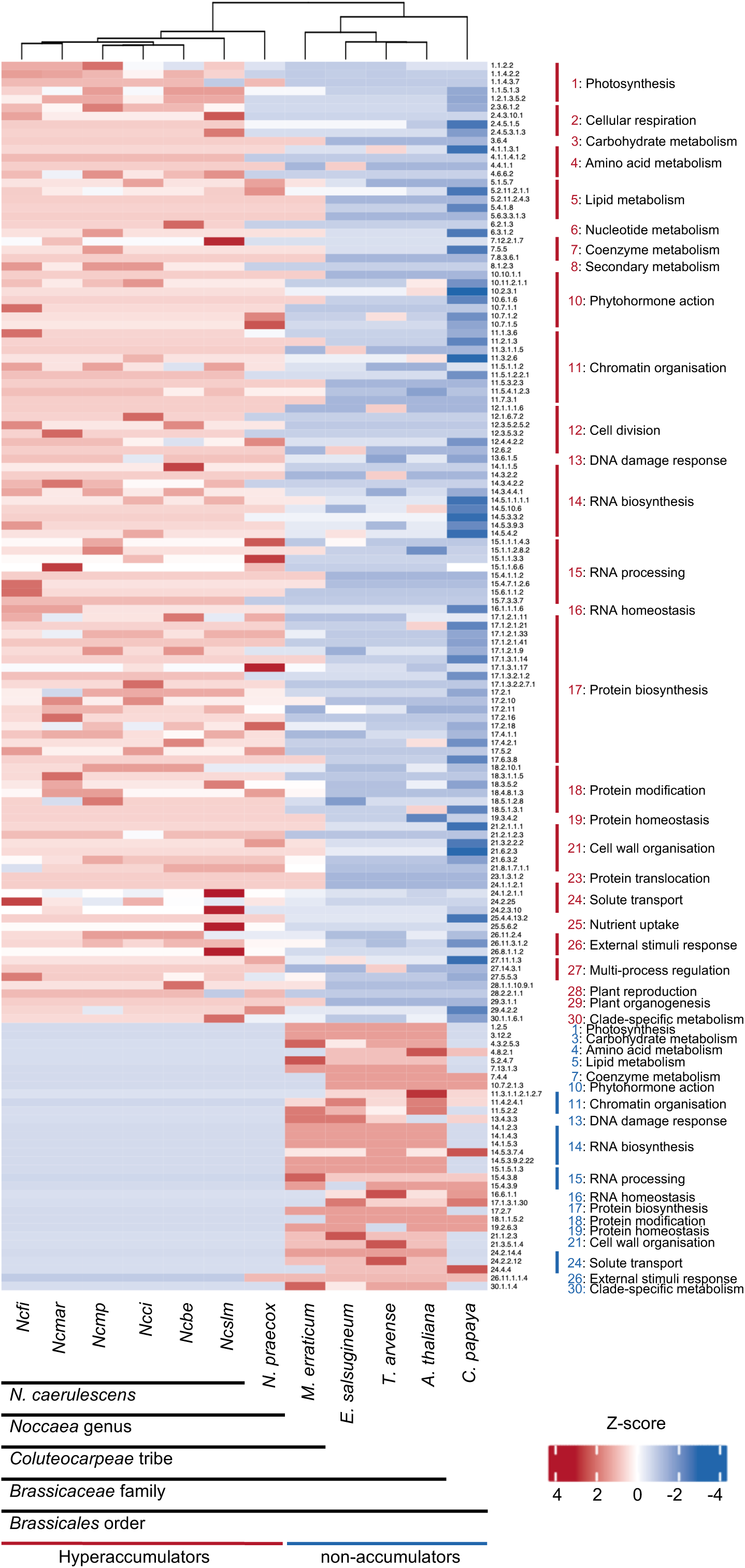
Gene copy number variation between metal hyperaccumulating and non-accumulating species of the Brassicaceae family. The heatmap displays the variation in gene copy number between metal hyperaccumulating species of the *Noccaea* genus (*N. caerulescens* and *N. praecox*), and non-accumulating species of the Brassicaceae family. *Carica papaya* of the Brassicales order is used as an outgroup. The genes are categorized according to MapMan functional bins and sub-bins. In each sub-bin, gene copy number is normalized across the species using a z-score. Only sub-bins showing a significant difference (Kruskal-Wallis test, p<0.001) between the hyperaccumulator and non-accumulator groups are displayed.

**Fig. 4:**
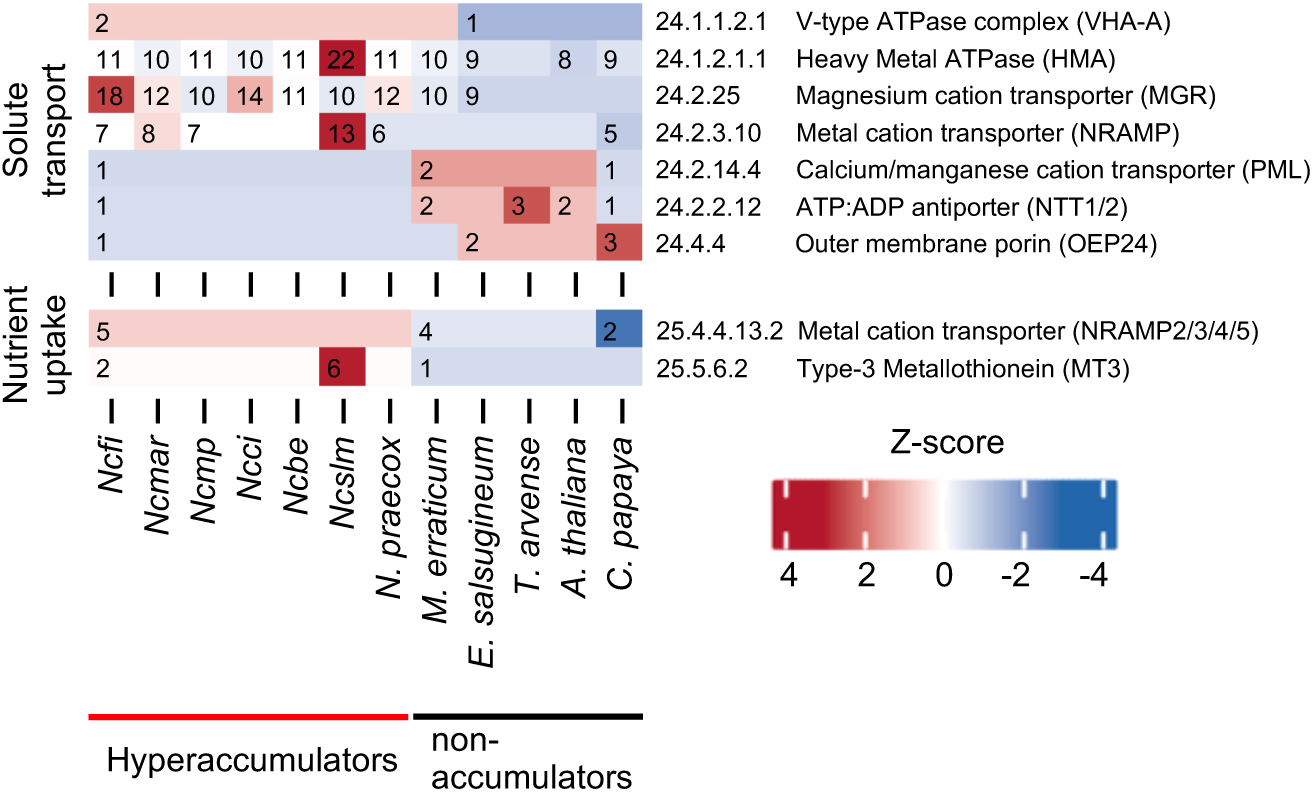
Gene copy number variation in genes involved in solute transport and nutrient uptake. The heatmaps display the variation of gene copy number between metal hyperaccumulating and non-accumulating species of the Brassicaceae family for genes involved in solute transport (MapMan bin 24) and nutrient uptake (MapMan bin 25). Gene numbers and standard scores (z-score) are displayed for each sub-bins. Only sub-bins showing a significant difference (Kruskal-Wallis test, p<0.001) between the hyperaccumulator and non-accumulator groups are displayed.

### 3.4. Analysis of gene copy number variation in gene families involved in metal transport and homeostasis

We next analyzed more specifically CNV in gene families coding for proteins involved in solute transport and nutrient uptake. The analysis of genes coding for Heavy Metal ATPases (HMAs, sub-bin 24.1.2.1.1) revealed that seven out of the eight *HMA* genes present in *A. thaliana* exhibit CNV among species of the Brassicaceae family (Fig. 5A). Notably, *NcHMA4* exhibits at least one tandem gene duplication on chromosome NC3 in all *N. caerulescens* accessions and in *N. praecox*, while it is a single gene in other non-accumulating Brassicaceae species including *M. erraticum* (Fig. 5B). The analysis of the genomic context around the *NcHMA4* duplication indicates that it is associated with the insertion of MITE elements upstream of the coding sequence that are absent in *M. erraticum* (Fig. 5B, Fig. 6A). All *N. caerulescens* accessions, except *Ncmar*, also contain a distal duplication of *HMA3* on chromosome NC2 in addition to the ancestral *HMA3* copy located next to *HMA2* on chromosome NC7. This distal duplication of *HMA3* is absent in *N. praecox* and in the other Brassicaceae species. The *N. caerulescens Ncslm* accession is of particular interest because it exhibits four tandem duplications of *HMA4* (Fig. 6A), as well as five tandem duplications of the *HMA2/HMA3* locus (Supplemental Fig. 6A). However, the duplicated *HMA2* copies are truncated and likely not functional.

**Fig. 5:**
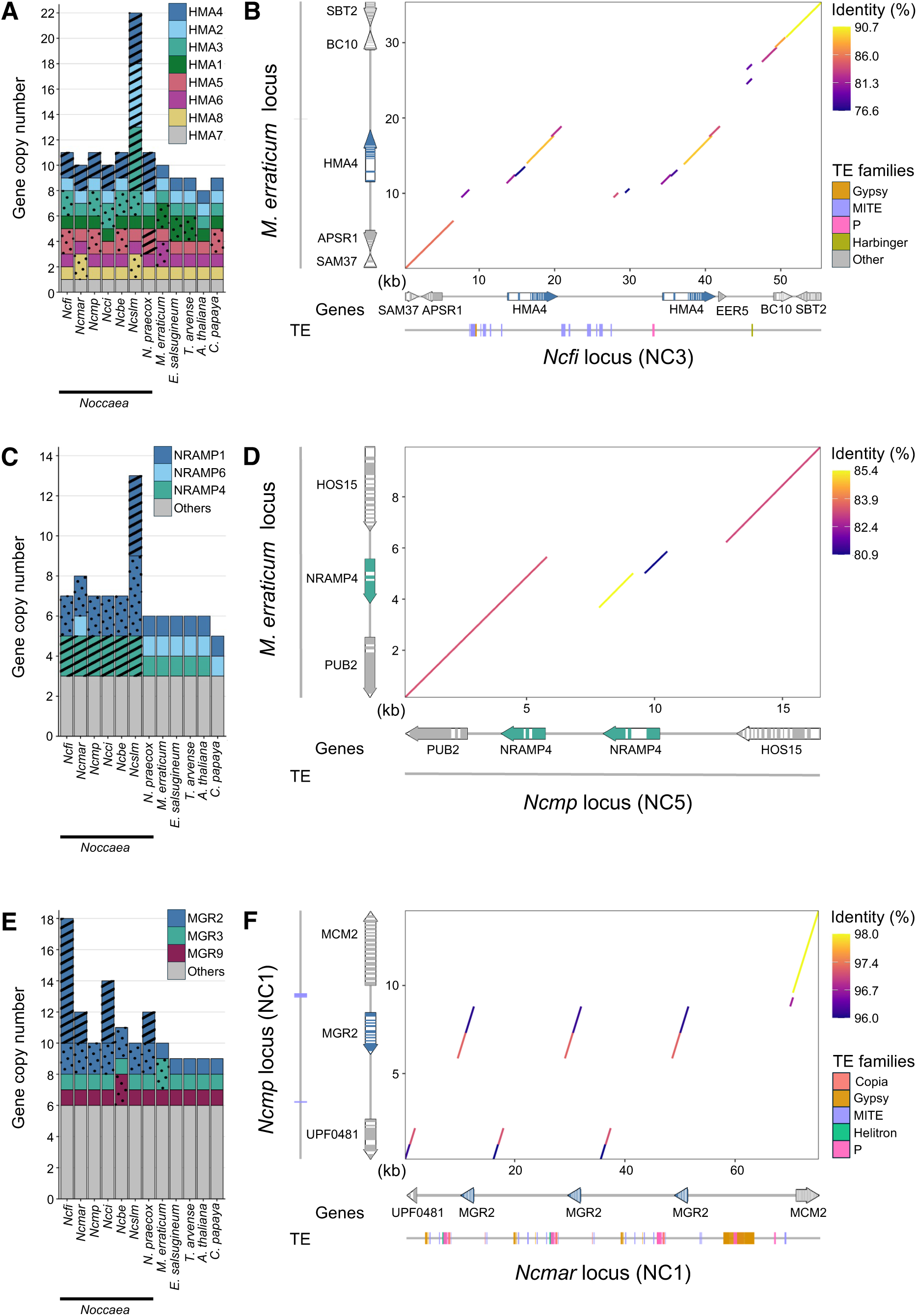
Gene copy number variation and genomic organization of selected metal homeostasis gene families in *Noccaea caerulescens* accessions. (A) Barplot showing gene copy number variation (CNV) for genes encoding HMA transporters across *Noccaea* species and *non-*accumulating Brassicaceae species. Gene duplications are classified as local (tandem) duplication (hatch pattern), or dispersed duplication (spot pattern). (B) Representation of the *HMA4* genomic locus in *Microthlaspi erraticum* and *Ncfi* (NC3) showing the positions of annotated genes and transposable elements (TEs). The dot plot display pairwise nucleotide sequence similarity between the selected loci. The alignment blocks are represented by segments connecting homologous coordinates between the query and reference sequences and are colored according to sequence identity (%). (C) Barplot showing CNV for genes encoding NRAMP transporters. CNVs were only observed for *NRAMP1*, *NRAMP4* and *NRAMP6*. (D) Representation of the *NRAMP4* loci in *M. erraticum* and *Ncmp* (NC2) illustrating the duplication of this gene in *N. caerulescens* accessions. (E) Barplot showing CNV for genes encoding MGR transporters. CNVs were observed for *MGR2*, *MGR3* and *MGR9* orthologs. (F) Representation of the *MGR2* locus on NC1 illustrating the tandem triplication of *MGR2* in *Ncmar* while this gene is present as a single copy in *Ncmp* at this locus.

**Fig. 6:**
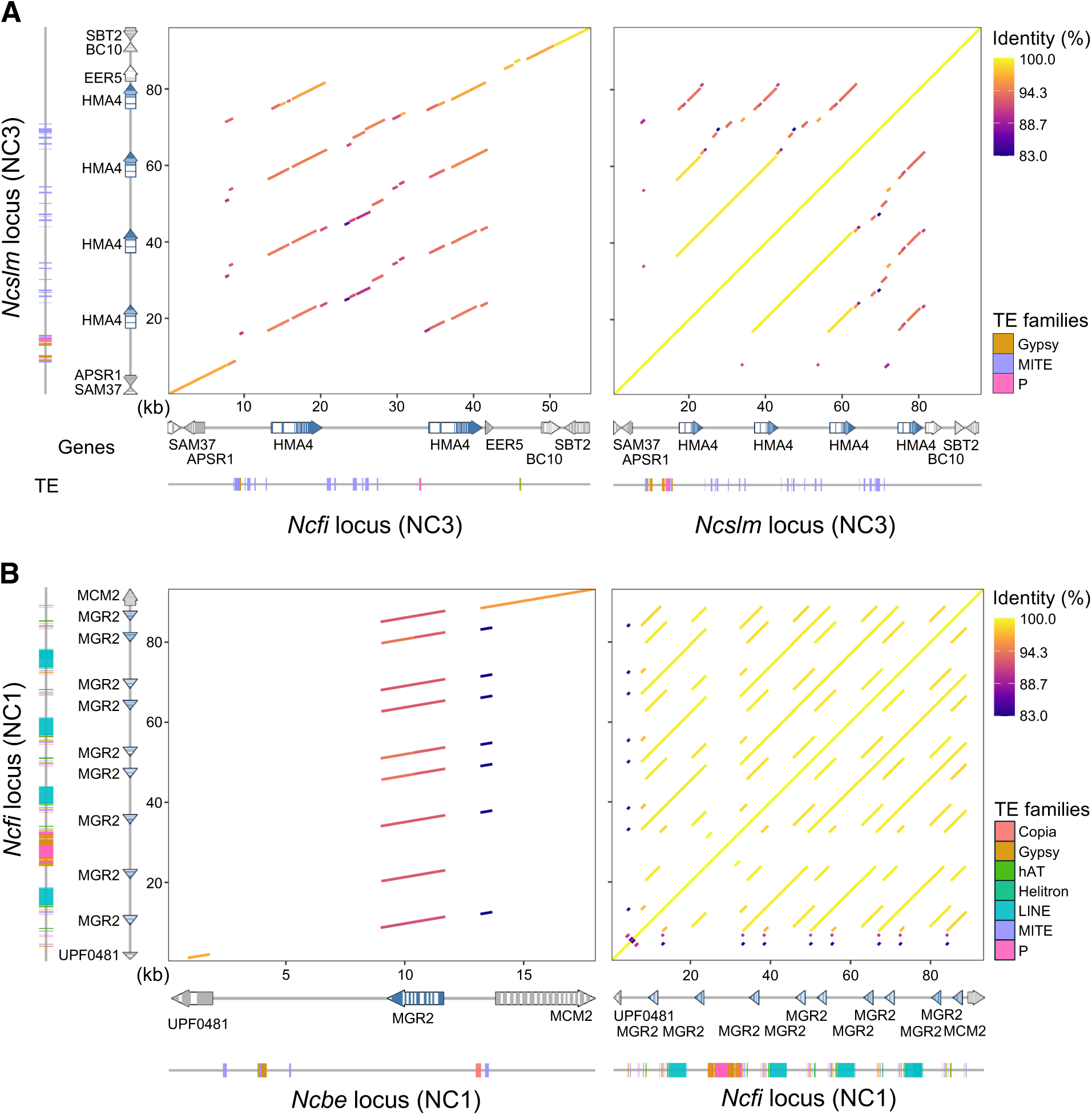
Recent expansion of *HMA4* and *MGR2* genes in *Noccaea caerulescens*. (A) Dot plots showing pairwise nucleotide sequence similarity at the *HMA4* locus on NC3 from *Ncslm* and *Ncfi* accessions. Alignment blocks are represented by segments connecting homologous coordinates between the sequences and are colored according to nucleotide sequence identity (%). The annotation of genes and transposable elements (TEs) are aligned with the dot plots. (B) Same representation as in (A) at the *MGR2* locus on NC1 from *Ncfi* and *Ncbe* accessions.

Our analysis also revealed gene CNV in the NRAMP transporter family (Fig. 5C). The *NRAMP4* gene is duplicated in tandem on chromosome NC5 in all *N. caerulescens* accessions, but not in the other species of Brassicaceae family including *N. praecox* and *M. erraticum* (Fig. 5D). We also observed a distant duplication of the *NRAMP1* gene on chromosome NC2, located approximately 160 kb away from the ancestral copy, in all *N. caerulescens* accessions (Fig. 5C). In the *Ncslm* accession, this distal copy underwent local recombination (Supplemental Fig. 6B). Overall, the *Ncslm* accession has eight copies of *NRAMP1*, originating from both additional distal duplications on chromosomes NC1 and NC6, and from tandem duplications at the locus on NC1 (Supplemental Fig. 6B). In contrast, the *NRAMP6* gene, which is closely related to *NRAMP1*, and present as a single-copy gene in Brassicaceae species, is apparently absent in *N. caerulescens* except in the *Ncmar* accession (Fig. 5C).

More surprisingly, our analysis revealed CNV for genes encoding magnesium cation transporters of the MGR family (Fig. 5E). In addition to the conserved copy of *MGR2* associated with *MGR1* in all Brassicaceae species (Tang et al., 2022), and present on chromosome NC7 in *N. caerulescens*, *Noccaea* species contain a second copy of this gene on chromosome NC1 (Fig. 5F). This distant copy is further duplicated in tandem from two to eight copies in the *N. caerulescens* accessions *Ncmar*, *Ncci* and *Ncfi*, as well as in *N. praecox*. These later duplications are associated with distinct TE landscape in these accessions suggesting independent TE insertion events. (Fig. 5F, Fig 6B).

### 3.5 Duplications affecting other genes involved in metal homeostasis

While genes encoding metal transporters of the ZIP family have been implicated in metal hyperaccumulation in *N. caerulescens* (Belloeil et al., 2026; Halimaa et al., 2019, 2014), our comparative analysis did not reveal significant change in gene CNV for this class of transporters (sub-bins 24.2.11.1 and 25.4.1.1.2.1) between the hyperaccumulator and the non-accumulator groups. However, we observed gene duplication events in the IRT1-clade of ZIP transporters in species of the Coluteocarpeae tribe, including *M. perfoliatum*, that are absent from other Brassicaceae species (Fig. 7A). The *IRT2* gene is duplicated in tandem in all species of the Coluteocarpeae tribe. Two distant copies of *ZIP7* are present in all *N. caerulescens* accessions while it is a single copy gene in *N. praecox* and *M. erraticum*. We also detected tandem or distant duplication of *ZIP10* in *N. caerulescens* accessions *Ncmar* and *Ncmp* as well as in *N. praecox*. Finally, we identified a tandem duplication of IRT1 specific to the *Ncfi* accession, associated with the insertion of a Gipsy LTR-retrotransposon (Fig. 7C).

**Figure 7.**
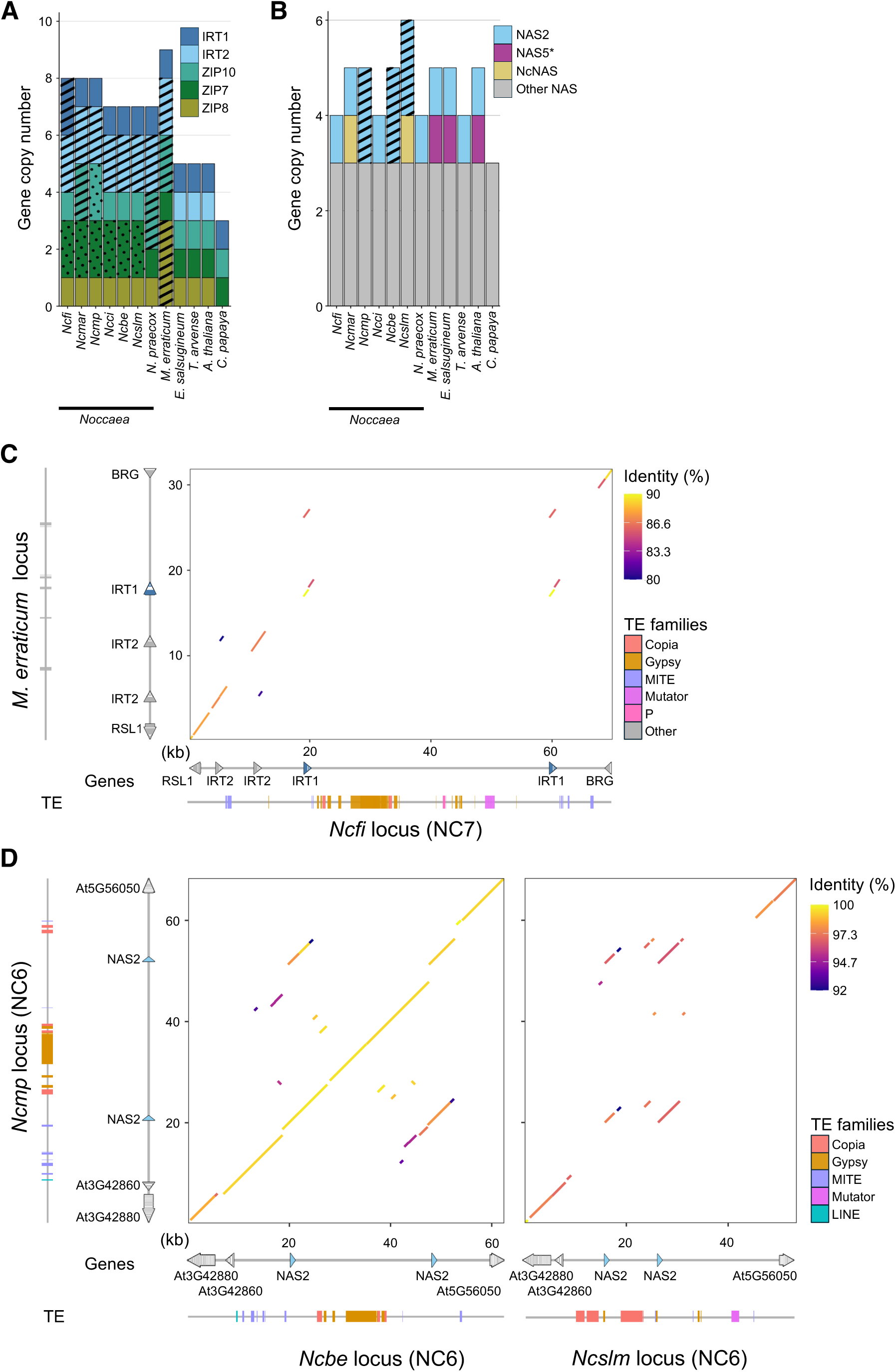
Copy number variation for *ZIP/IRT* and *NAS* gene families in *Noccaea caerulescens*. (A) Barplot showing CNV for *ZIP/IRT* family genes (IRT1 clade) in Brassicaceae species. (B) Barplot showing gene copy numbers for *NAS* family genes. *Several Brassicaceae species contain a *NAS5* gene, including *Arabidopsis halleri*, annotated as a pseudogene in *A. thaliana* (Seebach et al., 2023). NcNAS is a clade of *NAS* genes identified in some *N. caerulescens accessions*. (C) Representation of the *IRT1* genomic locus in *Microthlaspi erraticum* and *Ncfi* (NC7) showing the positions of annotated genes and transposable elements (TEs). The dot plot display pairwise nucleotide sequence similarity between the selected loci. The alignment blocks are represented by segments connecting homologous coordinates between the query and reference sequences and are colored according to sequence identity (%). (D) Representation of the *NAS2* genomic locus (NC6) in *N. caerulescens* accessions *Ncmp*, *Ncbe* and *Ncslm*.

The high expression of nicotianamine synthase genes (*NAS*), which are responsible for synthesizing the metal chelator nicotianamine, has been associated with metal hyperaccumulation (Cornu et al., 2015; Halimaa et al., 2014). We identified a tandem duplication of *NcNAS2*, in the accessions *Ncmp*, *Ncbe* and *Ncslm* (Fig. 7B), that is associated with distinct TE landscapes in these accessions (Fig. 7D). In addition, we observed two *NAS* genes, called *NcNAS*, in the *Ncmar* and *Ncslm* accessions, that diverged from the other *NAS* genes identified in Brassicaceae species (Fig. 7B).

These results indicate that additional CNV affecting genes involved in metal homeostasis were not identified in our comparative analysis because they are more generally present in the Coluteocarpea tribe or present in only few *N. caerulescens* accessions. These duplications may have favored the evolution of metal hyperaccumulation in *Noccaea* species or are involved in local adaptation, respectively.

## 4. Discussion

The primary objective of this study was to expand our knowledge on the genomic diversity of the model metal hyperaccumulator species *N. caerulescens*. Here, we demonstrate that using Oxford Nanopore Technologies (ONT) long reads alone or together with PacBio HiFi reads significantly improved the quality of the *N. caerulescens* assemblies compared to the previous *Ncci* v2.3 assembly generated with PacBio HiFi reads and scaffolded with Hi-C chromosome conformation capture data (Belloeil et al., 2026). The hybrid *Ncci* v3.1 assembly directly generated longer contigs, including those that fully cover chromosomes NC1, NC3, and NC7 (Fig. 1; Table 1). The assembly of the genomes of the other *N. caerulescens* accessions further indicated that assembling ONT reads at a coverage above 70X resulted in a quality of assembly similar to that of *Ncci* v3.1. Conversely, the assembly of PacBio HiFi reads from *Ncslm* generated smaller contigs (Table 1). Obtaining large contigs greatly facilitates chromosome scaffolding. The comparison of the *Ncci* v3.1 and *Ncci* v2.3 assemblies revealed that two large inversions had been previously introduced in chromosomes NC3 and NC5 (Fig. 1B) likely due to the presence of long-stretch of repetitive sequences in the genome of *N. caerulescens* that affected the interpretation of Hi-C data during the scaffolding of the *Ncci* v2.3 assembly. The *Ncci* v3.1 assembly conforms more closely to the previously proposed structure of the *N. caerulescens* genome based on cytogenetic analysis (Mandáková et al., 2015). We used the *Ncci* v3.1 assembly as a reference for the scaffolding of chromosomes in the other *N. caerulescens* accessions. We did not observe any conflicts between the scaffolded chromosomes and the linear sequence of the contigs. This indicates that the scaffolding step did not overwrite the structure deduced by directly assembling long ONT reads. Furthermore, despite using the *Ncci* v3.1 assembly as a reference, we observed large inversions in the lower arm of chromosome NC2 and in the upper arm of chromosome NC5 that are specific to the *Ncci* and *Ncmar* accessions respectively (Fig. 2). Apart from these inversions, the overall structure of the *N. caerulescens* genome appears to be conserved in *N. caerulescens* accessions from different genetic origins (Gonneau et al., 2017; Mandáková et al., 2015).

Chromosome-quality assemblies for several accessions of *N. caerulescens* allowed us to conduct genomic comparative analysis and identify gene copy number variation (CNV) associated with metal hyperaccumulation within the *Noccaea* genus. Gene duplication is an important driver of plant evolution and adaptation as it can result in increased gene expression and provide opportunities for gene neofunctionalization (Panchy et al., 2016). The high expression of genes encoding the metal transporters *HMA3*, *HMA4* and *NRAMP1* has been associated with gene duplication in *N. caerulescens* (Craciun et al., 2012; Milner et al., 2014; Ó Lochlainn et al., 2011; Ueno et al., 2011). Our analysis of CNV confirmed the expansion of the *HMA4* and *NRAMP1* genes, and reveal the duplication of the magnesium transporter gene *MGR2* in all *N. caerulescens* accessions (Fig. 4, Fig. 5). However, our analysis also indicates that gene CNV affects a wide variety of biological functions in *N. caerulescens*, and that genes involved in metal homeostasis, which are essentially present in MapMan bins 24 and 25, are not over-represented among all those affected by CNV (Fig. 3). Further work will be required to establish whether other functions affected by CNV are related or independent of the evolution of metal hyperaccumulation in the *Noccaea* genus.

All *N. caerulescens* accessions contain at least a tandem duplication of the *HMA4* gene, which is not observed in non-accumulating Brassicaceae species. The number of *HMA4* duplications is higher in *Ncslm* (four duplications) and in *N. praecox* (three duplications) (Fig. 5A, Fig. 6A). *N. caerulescens* accessions also contain a distant *HMA3* duplication on chromosome NC2, in addition to the ancestral *HMA3* copy associated with *HMA2* on chromosome NC7.

This distant duplication of *HMA3* is absent in the *N. caerulescens* accessions *Ncmar* and in *N. praecox*. Conversely, *HMA3* has undergone further tandem duplications, reaching seven copies in *Ncslm,* as previously observed in the related Ganges accession (Ueno et al., 2011). This result suggests that the high copy number of *HMA3*, which is involved in the sequestration of cadmium in the vacuole (Morel et al., 2009; Ueno et al., 2011), is fixed in the *N. caerulescens* populations originating from calamine soils in the southern part of Massif Central in France. This provides a genetic support for these populations’ specific capacity to accumulate and tolerate high cadmium concentrations (Sterckeman et al., 2017). Our results also reveal that all *N. caerulescens* accessions have a tandem duplication of *NRAMP4* and a distant duplication of *NRAMP1* absent from *N. praecox*. The *Ncslm* accession stands out once again with eight copies of *NRAMP1* originating from distant and tandem duplication of this locus. These results confirm and refine previous findings on *NcNRAMP1* CNV in *N. caerulescens* accessions, as determined by quantitative PCR (Milner et al., 2014) . They also further support the role of this transporter in cadmium uptake in *N. caerulescens* (Halimaa et al., 2014; Milner et al., 2014). The effect of the duplication of the vacuolar export transporter *NcNRAMP4* on metal hyperaccumulation remains unclear, though it could be involved in tolerance to metals and oxidative stress (Molins et al., 2013; Oomen et al., 2009).

More surprisingly, our analysis revealed gene expansion in the sub-bin 24.2.25, which corresponds to the metal transporters of the Ancient Conserved Domain Proteins (ACDP)/CNNM family, also known as Magnesium Release Transporters (MGRs) in plants (Tang et al., 2022). Specifically, all of the *Noccaea* genomes that were investigated here contain a distant duplication of *MGR2*, which is absent in non-accumulating species of the Brassicaceae family (Fig. 6C). This distant copy underwent additional independant tandem duplication in the *N. caerulescens* accessions *Ncfi*, *Ncci* and *Ncmar*, and in *N. praecox* (Fig. 5C, Fig. 6B). MGR2 was recently described in *A. thaliana* as a transporter that mediates magnesium transport in the vacuole and was proposed to play a role in adaptation to magnesium excess (Tang et al., 2022). Thus, the duplication of *MGR2* in *Noccaea* species may be indicative of an ancestral adaptation to ultramafic soils, which are characterized by a high Mg/Ca imbalance. Alternatively, the duplication of *MGR2* could be directly linked to metal accumulation, as ACDP/CNNM transporters can display a broad specificity for divalent metals (Chen and Gehring, 2023; Goytain and Quamme, 2005). In contrast to these genes, the duplication of genes that encode metal transporters of the ZIP family do not appear to be as strongly correlated with the metal hyperaccumulation trait in *Noccaea* species. We observed an increased copy number of *ZIP* genes of the IRT1-clade, including *IRT2* and *ZIP10* (Mäser et al., 2001), in several *Noccaea* species, as well as in the non-accumulator *Microthlaspi erraticum* of the Coluteocarpea tribe (Fig. 7A). The biological significance of *ZIP* genes duplication in this non-accumulating species is unclear, but it may be associated with the adaptation to calcareous soils, in which metals such as iron are poorly available (Mishra et al., 2020). On the other hand, the duplication of *ZIP* genes to acquire more metals in the common ancestor of the *Noccaea* and *Microthlaspi* genera may have favored the evolution of metal hyperaccumulation in *Noccaea* species. Interestingly, a tandem duplication of *NcIRT1* was specifically identified in the ultramafic accession *Ncfi* (Fig. 7C), consistent with the role of this transporter in nickel uptake and hyperaccumulation in *N. caerulescens* (Belloeil et al., 2026).

The duplication of genes involved in regulating metal homeostasis such as *HMA3*, *HMA4*, *NRAMP1*, *MGR2*, and *NAS2* in *N. caerulescens* correlates with their high expression in this species (Belloeil et al., 2026; Halimaa et al., 2014; Hammond et al., 2006; van de Mortel et al., 2006). However, CNV alone cannot explain the significant change in gene expression. For instance, *HMA4* was found to be expressed 30 times more in the roots of *N. caerulescens* accessions *Ncfi*, *Ncbe* and *Ncmp* than in *Microthlapi perfoliatum* (Belloeil et al., 2026), despite the identification of only two copies of *HMA4* in these accessions. Therefore, the duplication of these genes is also likely associated with the modification of their promoter, leading to strong cis-activation. In the hyperaccumulator *Arabidopsis halleri*, the high expression of *AhHMA4* results from gene triplication and the insertion of cis-acting elements (Hanikenne et al., 2008). Recently, it was discovered that the insertion into the promoter region of *AhHMA4* of Metal Hyperaccumulation Elements (MHE), responsible for the recruitment of the circadian transcription factor CCA1, was responsible for the high expression of this metal pump (Castanedo et al., 2025). This study also suggested that the presence of a similar motif in the promoter of the four copies of *HMA4* in *N. caerulescens* Ganges accession. However, we could not identify a conserved insertion of these MHE motifs in the promoters of *NcHMA4* copies present in other *N. caerulescens* accessions. Alternatively, we observed the insertion of miniature inverted-repeat transposable elements (MITEs) in upstream sequences of *NcHMA4* copies, which may have introduced cis-elements increasing the expression of this metal transporter (Soundiramourtty and Mirouze, 2025). Analyzing the promoters of the *NcHMA4* copies would allow to directly link the presence of MITEs to the high activity of these promoters (Iqbal et al., 2013). Surprisingly, we did not observe CNV at the *NcIREG2* locus, which encodes for a metal transporter involved in nickel hyperaccumulation, highly expressed in *N. caerulescens* and other nickel hyperaccumulators (Belloeil et al., 2026; García de la Torre et al., 2021). Therefore, we propose that the high expression of *IREG2* in *N. caerulescens* results from the insertion in its promoter of cis-regulatory elements that have yet to be identified. Recently, two *IREG2* genes have been identified in the nickel hyperaccumulator *Odontarrhena chalcidica* of the Brassicaceae family, suggesting that the duplication of *IREG2* may also be involved in the evolution of nickel hyperaccumulation in other species (Cai et al., 2026). Future research is also required to determine whether the evolution of metal hyperaccumulation involves neofunctionalization of duplicated genes.

Our data suggest that transposable elements (TEs) have play a role in *N. caerulescens* adaptation to various type of soil and in the evolution of metal hyperaccumulation. TEs are known to play an important role in genome evolution and adaptation mechanisms by favoring genome rearrangement, gene duplication, and the insertion of cis-elements, as well as causing gene disruption (Dubin et al., 2018; Lisch, 2013). In this study, we confirmed a high proportion of repeated sequences and TEs in the genome of six accessions of *N. caerulescens* (Table 1; Supplemental Fig. 3). In addition to the insertion of MITEs elements upstream of *NcHMA4* copies, our analyses indicate that Class I TEs are inserted in the vicinity genes of duplicated in *N. caerulescens*. Long terminal repeat (LTR) retrotransposons of the Gypsy and Copia families are associated with the duplication of *NcIRT1* in *Ncfi*, and with the duplication of *NcNAS2* in *Ncbe*, *Ncmp* and *Ncslm* (Fig. 7C and D). Conversely, the nine tandem copies of *NcMRG2* observed in *Ncfi* are associated with long interspersed nuclear elements (LINE) retrotransposon. The high sequence identity in the different *NcMRG2* coding sequences and the LINE retrotransposon copies suggests that this duplication burst originated from a recent duplication event (Fig. 6B). It has been proposed that LTR and LINE retrotransposons can stimulate tandem duplication by providing sources of sequence homology that favor unequal crossing over or co-mobilize adjacent genes during transposition (Cerda-Hernandez and Bureau, 2026; Hu et al., 2022). However, the tandem duplications of the *NcHMA4* and *NcNRAMP4* loci are not visibly associated with retrotransposon insertions (Fig. 5B and D, Fig. 6A). Finally, the diversity of TEs identified at duplicated loci such as *MGR2* and *NAS2* in several accessions (Fig. 5F, Fig. 6B, Fig. 7C), suggest that the capacity to adapt to different edaphic conditions may have evolved independently but convergently in several accessions of *N. caerulescens*.

## Authors contribution statement

Maria Kuzmik: Investigation, Validation, Visualization, Writing - Original Draft. Ophélie Courtin: Software, Formal analysis, Data Curation, Visualization, Writing - Original Draft. Christophe Klopp: Methodology, Software, Supervision. Marie-Christine Carpentier: Software, Formal analysis. Nathalie Rodde: Investigation. Caroline Callot: Investigation. Anne Ausset: Investigation. Céline Roques: Investigation. Céline Vandecasteele: Investigation. Véronique Gautier: Investigation. Elodie Belmonte: Investigation. Elodie Flaven: Investigation. Céline Devaux: Resources, Supervision, Funding acquisition. Agnès Mignot: Investigation. Marie Mirouze: Validation, Supervision. Sylvain Merlot: Conceptualization, Validation, Writing - Review & Editing, Supervision, Project administration, Funding acquisition.

## Data Availability

The Python scripts used in this study can be found at https://github.com/courtin-ophelie. The raw sequencing reads and assembled genomes generated in this work are available at NCBI’s BioProject PRJNA1096268. New *N. caerulescens* genome assemblies and their annotations were deposited in the recherche.data.gouv repository (doi:10.57745/OIP19T)

## Acknowledgements

We thank Mark Aarts (WUR, The Netherland) for providing seeds of *N. caerulescens* accessions, and Arnaud Bellec (CNRGV, France) for his support with ONT long-read sequencing. We also thank Anne-Claire Cazalé (LIPME/LRSV, France) for careful reading of the manuscript. This work was performed in collaboration with the GeT core facility (Toulouse, France, doi.org/10.15454/1.5572370921303193E12). GeT core facility was supported by France Génomique National infrastructure, funded as part of “Investissement d’avenir” program managed by Agence Nationale pour la Recherche (ANR-10-INBS-09). This work was supported by the French Laboratory of Excellence project "TULIP" (ANR-10-LABX-41; ANR-11-IDEX-0002-02), the École Universitaire de Recherche TULIP-GS (ANR-18-EURE-0019) and the ANR funded project HOMONIME (ANR-25-CE20-4366).

---

**Supplemental Fig. 1:**
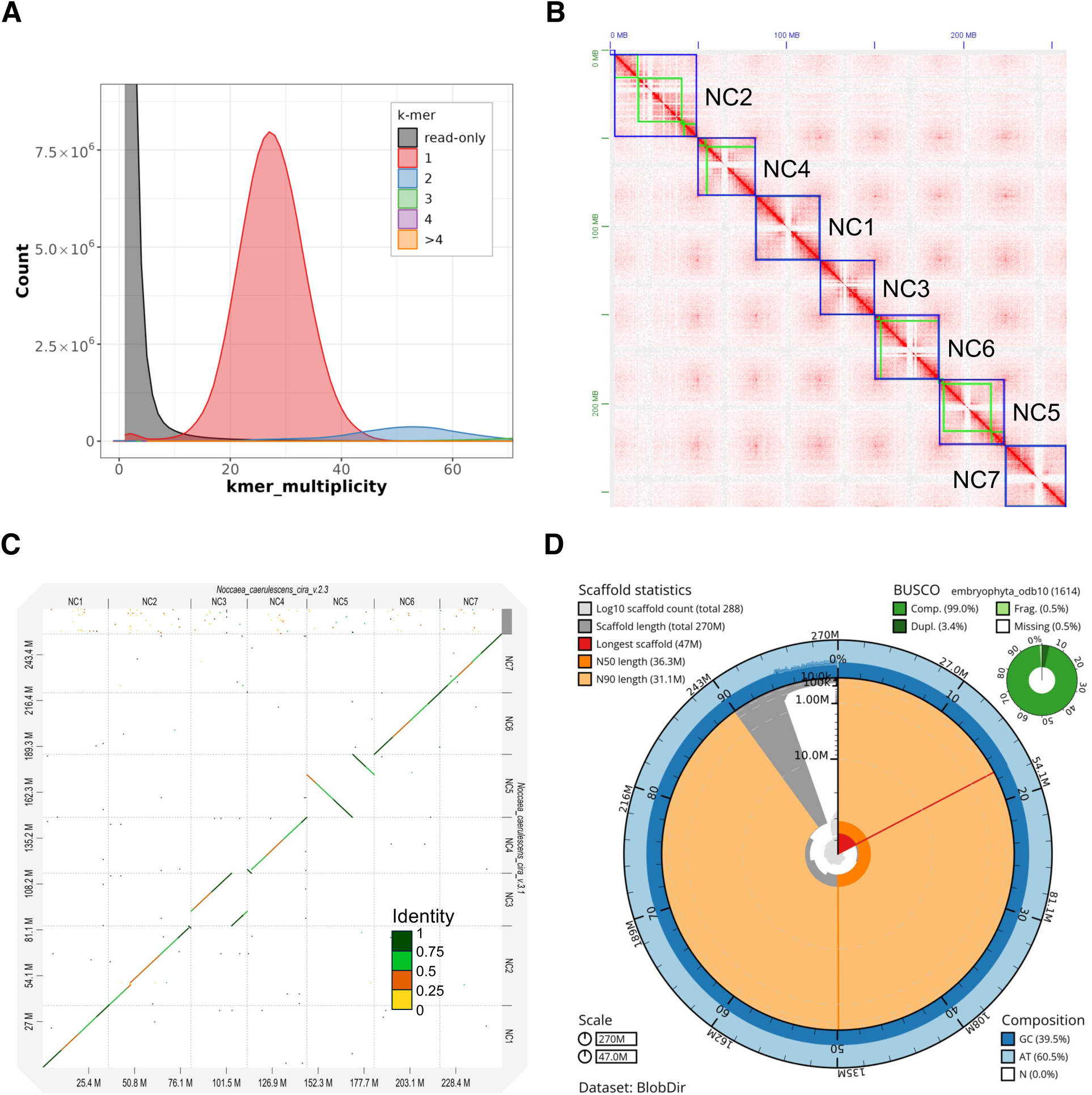
Quality assessment of the *Noccaea caerulescens* Cira genome assembly *Ncci* v3.1. (A) Merqury k-mer spectrum plot of the *Ncci* v3.1 assembly. (B) The Hi-C contact map was generated by remapping *N. caerulescens* Cira Hi-C reads to the *Ncci* v3.1 assembly. Green squares represent individual contigs. Blue squares represent the proposed seven chromosomes (NC1-NC7). (C) A D-GENIES comparison of the sequences of the seven chromosomes (NC1-NC7) proposed in the *Ncci* v2.3 and *Ncci* v3.1 assemblies. Alignment segments indicate homologous regions between the assemblies, with colors representing sequence identity. (D) A snail plot generated by BlobTools summarizing the *Ncci* v3.1 assembly metrics and completeness statistics.

**Supplemental Fig. 2:**
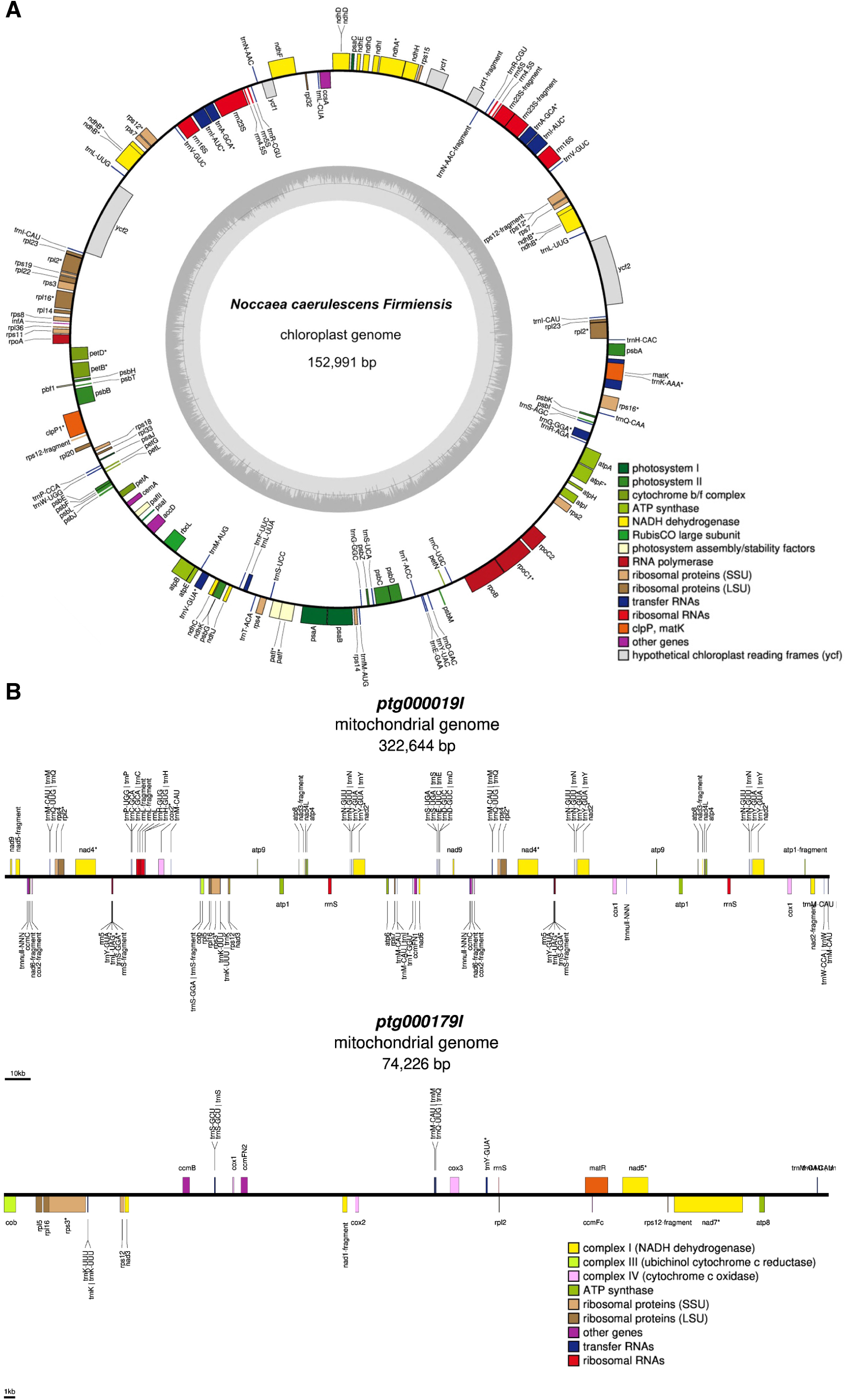
Chloroplast and mitochondrial genomes of *Noccaea caerulescens*. (A) Circular representation of the *Ncfi* chloroplast genome. Annotated genes are indicated on the outer ring, inside or outside depending on their transcription orientation, and are color-coded according to their functional. The inner track represents the variation in GC content along the chloroplast genome. (B) Linear representation of the *Ncfi* chloroplast mitochondrial genome. The *Ncfi* mitochondrial genome is composed of two contigs. Genes are displayed along the assembled genomic sequences and are color-coded according to their function.

**Supplemental Fig. 3:**
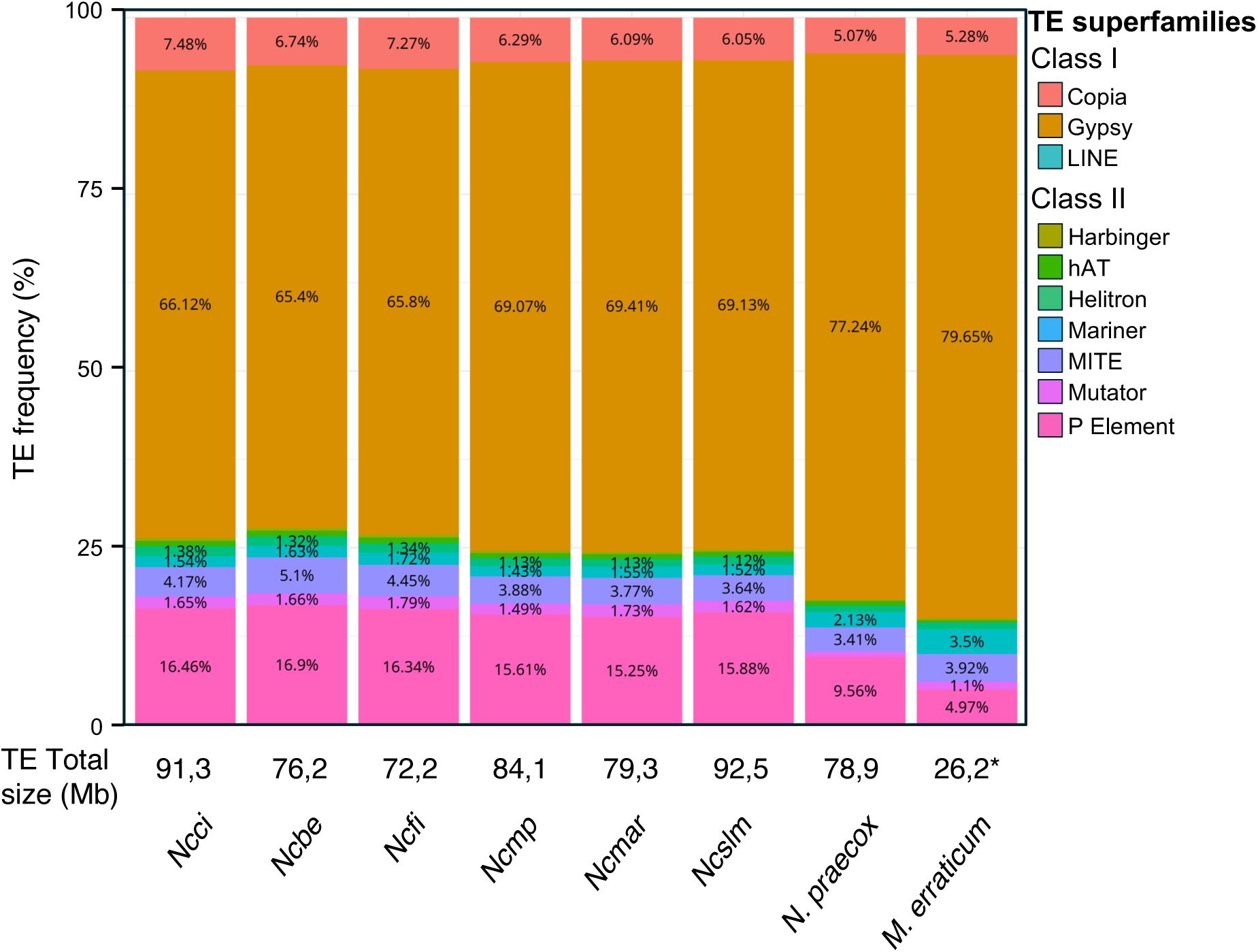
Annotation of transposable element (TE) superfamilies across *Noccaea caerulescens* accessions and related species of the Coluteocarpeae tribe. Bar plots showing the relative abundance of TE superfamilies in the six *N. caerulescens* accessions (*Ncci, Ncbe, Ncfi, Ncmp, Ncmar, and Ncslm*), as well as in *N. praecox* and *Microthlaspi erraticum*. TE superfamilies are divided into Class I (retrotransposons) and Class II (DNA transposons). The total size of TE in each genome is indicated. * Note that the *M. erraticum* genome assembly is more fragmented than those of the *Noccaea* species, which may have affected the TE annotation.

**Supplemental Fig. 4:**
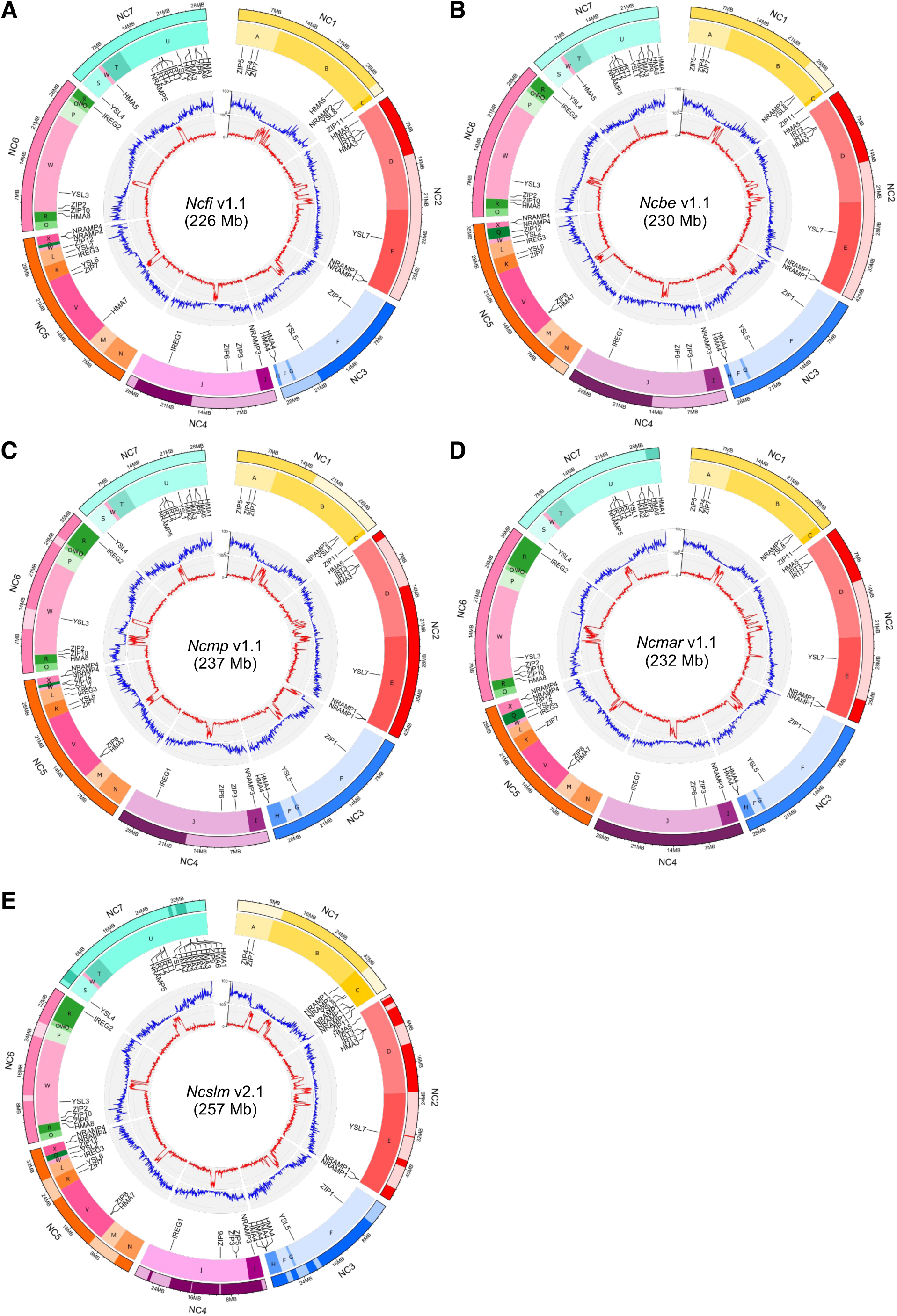
Representation of the genome assemblies of five *Noccaea caerulescens* accessions. Circos representation of the genome assemblies of (A) *Ncfi* v1.1, (B) *Ncbe* v1.1, (C) *Ncmp* v1.1, (D) *Ncmar* v1.1, and (E) *Ncslm* v2.1. The annotations are the same as in Fig. 1A.

**Supplemental Fig. 5:**
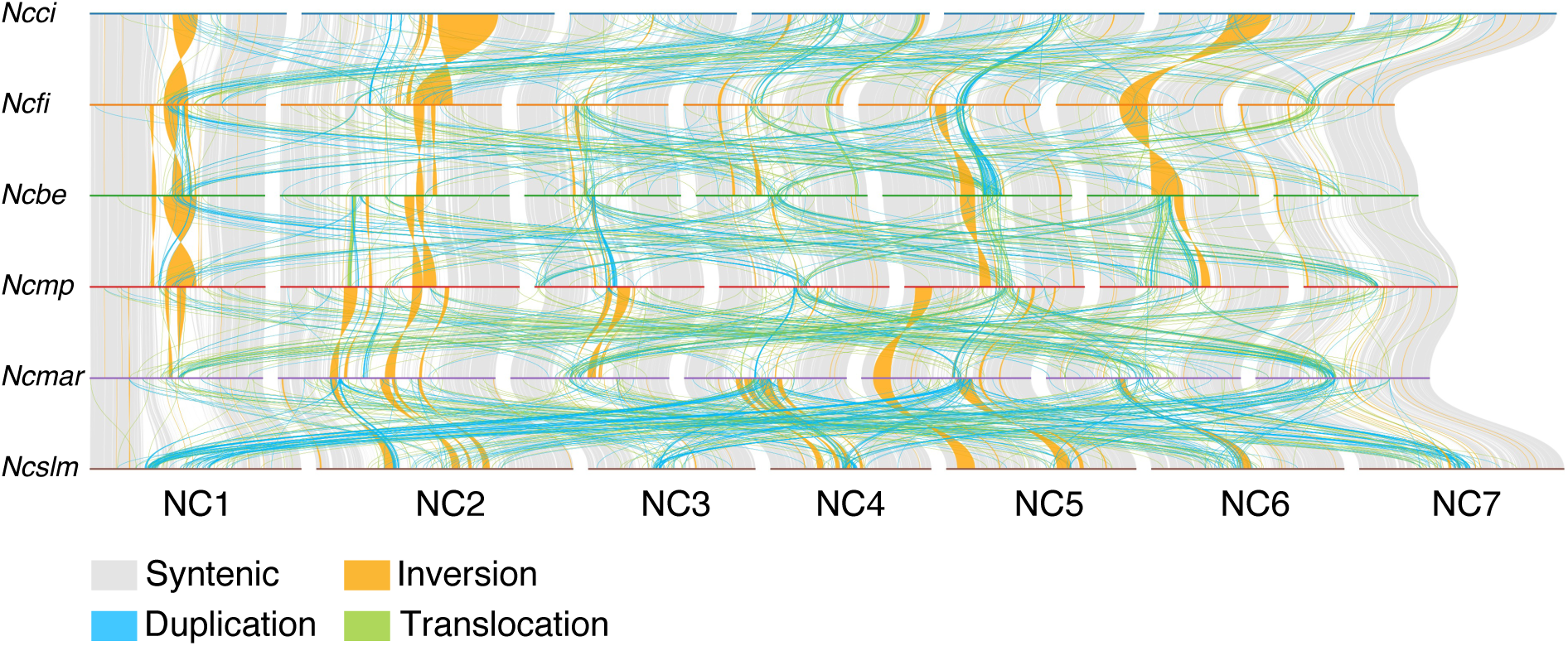
Comparison of the genome assembly of various *Noccaea caerulescens* accessions. The genome assemblies of the 6 *N. caerulescens* accessions described in this study are compared. The data are the same as in Fig. 2, but are organized by accessions to better visualize interchromosomal rearrangement. Syntenic regions and structural rearrangements, including duplications, inversions and translocations are indicated.

**Supplemental Fig. 6:**
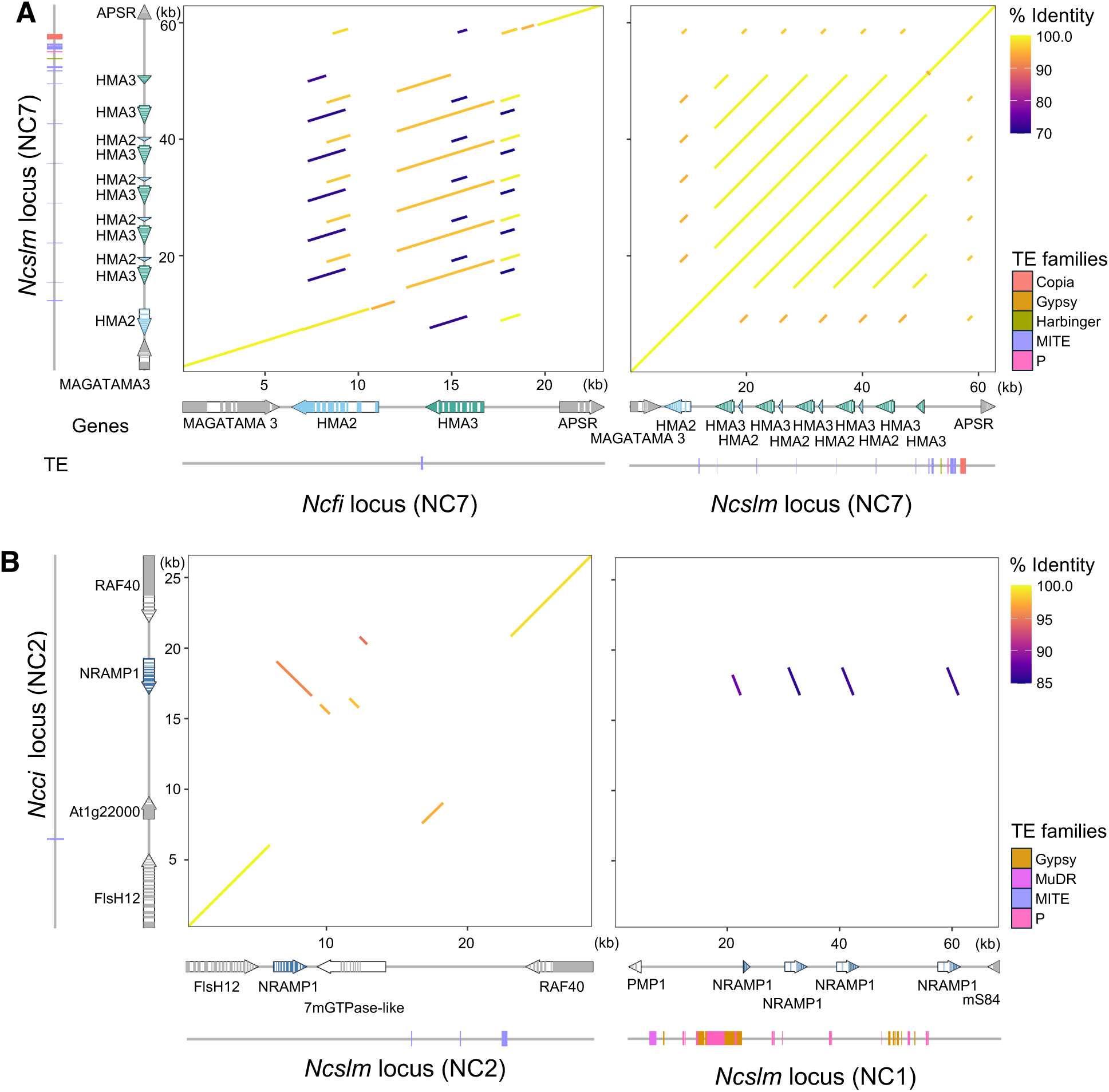
Organization of the *HMA2/HMA3* and *NRAMP1* loci in *Noccaea caerulescens*. (A) Representation of the *HMA2/HMA3* locus (NC7) in *Ncslm* and *Ncfi* showing the positions of annotated genes and transposable elements (TEs). The dot plot display pairwise nucleotide sequence similarity between the selected loci. The alignment blocks are represented by segments connecting homologous coordinates between the query and reference sequences and are colored according to sequence identity (%). (B) A comparative alignment of the *NRAMP1* locus on NC2 in *Ncci*, with the equivalent locus on NC2, as well with the distantly duplicated locus on NC1 in *Ncslm*.

