## Supplemental methods for "High-quality genome assembly of several *Noccaea caerulescens* accessions reveals gene expansion events associated with metal adaptation and hyperaccumulation"

*DNA extraction and sequencing*

For the *Ncci* accession*,* very high molecular weight genomic DNA was extracted from 6g of young frozen leaves using Qiagen Genomic-Tip G500 column (Qiagen, #10262) according to manufacturer instructions. Briefly, after grinding using liquid nitrogen, the fine powder is resuspended and briefly vortexed in 38ml of G2 buffer containing 76µl of 100mg/ml RNAse A (Macherey Nagel, #740505). After adding 2ml of 20mg/ml proteinase K (Applichem, #A3830.0100), the suspension in briefly vortexed and incubated at 50°C for 1hour under gentle horizontal rotation. After incubation, the lysate was centrifuged at 2000g for 20min at 15°C. The column was equilibrated using 10ml of QBT buffer (Qiagen, #19060). After centrifugation, the supernatant was transferred on the column and the filtration occurred by gravity. The column was washed twice with 15ml of QC buffer and the DNA eluted with 15ml of pre-heated (50°C) QF buffer. The DNA was precipitated using 0.7 volume of Isopropanol and the pellet collected by centrifugation (5000 g, 15 min, 4°C), washed with 70% ethanol, and then resuspended in EB buffer (Qiagen, # 19086). The quantity and purity of DNA were estimated using Qubit and Nanodrop (Thermofisher Scientific) and the fragment size was estimated using FEMTO pulse (Agilent). The DNA was sized using SageELF instrument (Sage Science) according to manufacturer instruction from 20 µg of DNA. Briefly, 20µg of DNA was loaded on 4 wells in Pre-Cast Agarose Gel Cassettes for DNA Fractionation (Sage Science, 5µg per well). The DNA migrates using the following program: Size-select 5-100Kb step3h (separate for 03:00:00 at 80.0 V with wave index 3-2, elute for 01:30:00 at 50.0 V with wave index 3-1 and finally rev elute for 00:00:05 at 25.0 V with wave index 3-1). The DNA was recovered after 45min and purified using 1X AMPure beads (Beckman Coulter). After DNA quantification and fragment size estimation, the fractions containing DNA fragments from 10 to 160 kb were pooled to prepare the library. The sequencing library was prepared using the ligation kit (Oxford Nanopore Technology, ref: SQK-LSK114) following the associated protocol (version GDE_9161_v114_revW_29Jun 2022) with minor changes. Briefy, 5µg of DNA was used and the reaction volume was multiplied by 4. The incubation times were increased to 30min and the final library was eluted in 15µl of the provided elution buffer. After quantification of the library and fragment size estimation, half of the library (962ng) was loaded onto a MinIon flow-cell (Oxford Nanopore Technology, ref: FLO-MIN114) following the manufacturer instructions. After 22h of run, the flow-cell was washed using the Flow Cell Wash Kit (Oxford Nanopore Technology, ref: EXP-WSH004) following the associated protocol (protocol version WFC_9120_v1_revQ _08Dec2020) and the remaining library was loaded for a second run. 5.4Gb of raw data was generated with a N50 of 38.61kb. The data were basecalled from pod5 files following the super-accurate model (sup) using Dorado tool (Oxford Nanopore Technology, v0.7.3, https://github.com/nanoporetech/dorado/).

For the *Ncslm* accession, high molecular weight DNA was extracted from 500 mg of frozen leaves that were ground with a pestle in a mortar cooled with liquid nitrogen. Four ml of pre-heated (50°C) extraction buffer (200 mM Tris-HCl pH8.0; 50 mM EDTA; 500 mM NaCl; 1.25 % SDS; 1% PVP 40000), containing 40 µl beta-mercaptoethanol, 200 µl CTAB (12,5%) and 10 µl RNase A (10mg/ml), were added to the leaf powder in a 50 ml tube and mixed by inversion and gentle vortexing. The mixture was incubated at 65°C for 20 min with gentle agitation and then placed on ice for 5 min. Five ml of ice-cold Potassium Acetate (3M/5M) was added to the tube and mixed by inversion before incubation on ice for 5 min. The mixture was centrifugated for 20 min at 4500 rpm using a swing-out rotor at 4°C. The supernatant was filtrated on Kimwipes paper towel and collected on a new 50 ml tube. The supernatant was splitted in 3 wells (3x1.5 ml) of a 24 deepwell plate for a KingFisher-Flex apparatus. In each sample, 30 µl of Chemagic magnetic beads (Perkin-Elmer, #CMG-252-A), 700 µl AMMCGE Buffer (33% Guanidium Chloride 7.8M, 67% Ethanol 96%) and 2 ml of isopropanol were successively added. The DNA was purified according to the KingFisher-Flex manufacturer protocol (INS-EXT-005-Kingfisher-Flex.pdf), and the Chemagen protocol (US patent 6 958 372 B2). The purified DNA (10 ug) was sheared to approximately 25 kb using a Diagenode Megaruptor (Diagenode) and DNA fragment size distribution verified using an Agilent Femto Pulse. The sheared DNA was used to prepare the library following the Procedure & Checklist - Preparing HiFi SMRTbell® Libraries using SMRTbell Express Template Prep Kit v2.0. The SMRTBell library was quality inspected and quantified on a Femto Pulse (Agilent Technologies) and a Qubit fluorimeter with Qubit dsDNA HS reagent Assay kit (Life Technologies). The ready-to-sequence SMRTBell Polymerase Complex was generated using a Sequel II Binding Kit 2 and the primer v2. The library (80 pM) was sequenced on a PacBio Sequel II instrument Sequel II Sequencing plate v2.0 (Pacific Biosciences), using CCS mode.

For the other *N. caerulescens* accessions, library preparation and sequencing were performed according to the manufacturer’s instructions “Ligation-sequencing-gdna-native-barcoding-v14-sqk-nbd114-24”. At each step, the DNA was quantified using the Qubit dsDNA HS Assay Kit (Life Technologies). DNA purity was tested using the nanodrop (Thermofisher), and the size distribution and degradation assessed using a Fragment Analyzer (Agilent) DNF-464 HS Large Fragment Kit. Purification steps were performed using AMPure XP beads (Beckman Coulter). For one Flowcell, 4 µg of each DNA sample (5 samples) were purified and then sheared at 30kb (speed V29) using a Megaruptor3 system (Diagenode). Using NBD114.24 kit (Oxford Nanopore Technologies), the library was generated by equimolar pooling of 1 µg of each barcoded sample. The library was loaded onto a R10.4.1 flowcell and sequenced on a PromethION instrument at 7 fmol within 72 hours.

*Identification of contigs corresponding to the chloroplast and mitochondrial genome*

The contigs corresponding to the chloroplast genome of the *N. caerulescens* accessions were identified by mapping to the *Thlaspi arvense* chloroplast genome (NC_034362.1) using minimap2 2.28 (Li, 2018). When multiple contigs were found to match the chloroplast genome, they were ordered and oriented based on the reference sequence, and subsequently concatenated into a single sequence by inserting stretches of 100 Ns between adjacent contigs to represent unresolved regions. The annotation and graphical representation of the chloroplast genome was performed with GeSeq ChloroBox (Tillich et al., 2017).

Mitochondrial contigs were identified using MitoHiFi 3.2.1 (Uliano-Silva et al., 2023) using the *Microthlaspi perfoliatum* mitochondrial genome as areference (OQ376512.1).
